# Metabolic starvation–induced cell swelling drives solid stress in tumors

**DOI:** 10.64898/2026.02.05.704098

**Authors:** Mohammad Dehghany, Vivek Sharma, Akash Samuel Annie-Mathew, Andrei Zakharov, Tom C. Hu, Guilherme Pedreira de Freitas Nader, Vivek B. Shenoy

## Abstract

Solid stress shapes tumor growth, invasion, and therapeutic response, yet its physical origin and clinical relevance remain unclear. Here, we develop a mechano–electro–osmotic model integrating metabolic gradients, ion transport, and cellular mechanics to explain residual solid stress emergence in tumor spheroids, common models of solid tumors. We show that solid stress arises predominantly from osmotic cell swelling driven by metabolic deprivation and ion accumulation, rather than proliferation. This mechanism generates a characteristic stress architecture: isotropic compression in the hypoxic core balanced by peripheral tangential tension, causing pronounced cell and nuclear deformation. The resulting nuclear strain provides a mechanical basis for DNA damage and genomic instability implicated in disease progression and treatment resistance. We validate these predictions in breast cancer using MDA-MB-231 spheroids and patient-derived ductal carcinoma in situ lesions, and corroborate them across published spheroid models and in vivo and ex vivo tumors spanning additional cancer types. Our findings link tumor metabolism to clinically relevant mechanical stresses, suggesting opportunities to target osmotic and metabolic pathways to mitigate solid stress and improve therapeutic outcomes.

## 1. Introduction

Abnormally elevated solid stresses are a hallmark of both benign and malignant tumors [1, 2]. These stresses originate from mechanical forces transmitted through the solid components of the tissue. A growing body of evidence demonstrates that elevated solid stresses profoundly influence tumor progression by altering cell behavior and undermining the effectiveness of therapeutic interventions. Specifically, solid stresses can promote cell migration [3, 4], enhance epithelial–mesenchymal transition (EMT) [3], and increase the invasive potential of cancer cells [5]. They also contribute to treatment resistance by inducing a quiescent state [6], thereby reducing tumor cell susceptibility to chemotherapies that target proliferating cells [6]. Moreover, elevated solid stresses can impair DNA repair, leading to genomic instability and increased intratumoral heterogeneity [7, 8], which facilitates the emergence of drug-resistant clones and enables tumors to evade treatment. As a result, strategies that relieve solid stress have shown promise in enhancing treatment efficacy [9-11]. Thus, a detailed mechanistic understanding of how solid stresses are generated within tumors is essential for guiding the development of more effective therapeutic interventions.

Solid stresses are thought to arise primarily from uncontrolled tumor growth and mechanical constraints imposed by the surrounding host tissue [2, 12, 13]. These stresses can be broadly classified into two components. One originates from reciprocal mechanical forces exerted by the surrounding normal tissue and is referred to as externally applied solid stress. The other is internally generated, stored within the tumor’s structural components, and persists even after the tumor is excised and external constraints are removed. This persistent component is known as growth-induced or residual solid stress [14]. While externally applied stress results from elastic compression of the host tissue, the mechanisms responsible for the generation of residual solid stress are more complex and remain an active area of research. Residual stresses can be estimated by making incisions in the tumor, allowing the stresses to relax, and measuring the resulting deformation at the site of the cut. Accordingly, a variety of cutting techniques, including partial cuts, planar incisions, tissue slicing, and needle biopsies, have been employed to assess residual stresses in animal and human tumors excised from various organs [15-18]. These experiments consistently reveal that residual stress in the core of most tumors is compressive and relatively isotropic, whereas stress at the periphery is predominantly anisotropic, a combination of circumferential tension and radial compression [13]. To investigate the origins of residual solid stress, researchers have also examined these stresses in multicellular tumor spheroids, which are three-dimensional in vitro cell aggregates that closely recapitulate key features of in vivo solid tumors [19, 20]. Upon reaching a critical size of approximately 500 µm, these spheroids accurately model avascular tumors and develop a stratified architecture comprising an outer proliferative layer, an intermediate zone of quiescent cells, and a central necrotic core that emerges due to steep gradients in oxygen and nutrient availability [21, 22]. Importantly, experiments have shown that partial incisions into spheroids consistently lead to outward protrusion of the central core (i.e., central bulging) and retraction of the peripheral layers [23-25], closely mirroring the deformations observed in tumors excised from various organs [15]. Together, these findings reveal a characteristic residual stress profile in the tumor tissues: central radial compression counterbalanced by tangential tension at the periphery. Yet, despite these extensive efforts, a mechanistic understanding of the origins of residual stress generation in tumor tissues remains lacking [13].

The stratified architecture of sufficiently large tumor spheroids and avascular tumors naturally gives rise to differential growth: proliferation is confined to the well-nourished periphery, while the interior becomes quiescent or necrotic due to limited oxygen and nutrients. This spatially heterogeneous growth is a well-established driver of residual stress in biological tissues [26-30], as localized expansion deforms the surrounding microstructure and generates internal stresses. Beyond proliferation, the volume of multicellular systems can also be regulated by swelling or shrinking of constituent cells, which in turn contributes to stress buildup. Cell volume is governed primarily by osmotic pressure, itself controlled by ion and water fluxes through channels, pumps, and gap junctions (GJs). In our earlier work [31], we demonstrated that in *small* tumor spheroids lacking notable metabolic gradients, intercellular solute flow through GJs drives peripheral swelling and central shrinkage. However, whether and how metabolic gradients in large spheroids and tumor tissues reshape spatial cell volume regulation, and how such gradients couple to residual solid stress generation, remains unknown.

Here, we address this open question by developing a mechano-electro-osmotic model that integrates metabolic gradients, cell proliferation, osmotic cell-volume regulation, and solid stress buildup in large tumor spheroids. We focus on spheroids because their well-defined spherical geometry and radially stratified organization provide a tractable, quantitative setting to study coupled transport, growth, and mechanics. Importantly, recent intravital measurements of solid stress at the cellular scale [32] have shown that stresses experienced by individual cells in tumor spheroids in vitro closely match those measured in vivo, establishing spheroids as a faithful physical model of tumor solid stress. Using our framework, we uncover a previously unrecognized mechanism of residual stress generation: impaired ion pump activity in nutrient-deprived cores drives osmotic swelling of central cells, which emerges as the dominant source of the characteristic compression–tension residual stress profile, surpassing the contribution of peripheral proliferation alone. The model further predicts a distinctive spatial organization of cell morphology, with rounded, swollen cells in the core and elongated cells in intermediate layers. We validate these predictions in breast cancer across complementary settings, from MDA-MB-231 spheroids to patient-derived human DCIS lesions, and establish generality by benchmarking against independent published datasets from other spheroid systems and tumor measurements obtained in vivo and after excision. Together, our results identify cell volume regulation as a central, previously overlooked driver of solid stress generation in tumors and establish a framework that can be extended to investigate how mechanical and metabolic cues jointly shape tumor growth and therapy response.

## 2. Development of a mechano-electro-osmotic model for solid stress in large tumor spheroids

To develop a theoretical framework that captures spatial variations in cell volume and their impact on residual solid stress generation in large tumor spheroids, we analyze local electrochemical and mechanical interactions between cells and the surrounding extracellular environment. These interactions regulate ion and water exchange, modulate cell volume, and give rise to macroscopic solid stress profiles by coupling system behavior across multiple scales. At the cellular level, mechanical and electrochemical interactions include active stresses generated by the actomyosin machinery, passive stresses arising from actin network deformation, and electrochemical forces generated by ion gradients and membrane potential. Together, these forces generate cortical tension and regulate cell shape and volume [33] (**Figure 1B**). Cells modulate their volume by exchanging water and ions through aquaporins, ion channels, pumps, and gap junctions (GJs) [34, 35] (**Figure 1C**). These exchanges depend on metabolic activity and the local availability of oxygen and nutrients. Experimental studies show that GJ conductance is highly sensitive to intracellular pH and decreases significantly when cytoplasmic pH falls into the high-6 range [36-40]. In large tumor spheroids, limited oxygen and nutrient availability lead to lactic acid accumulation [41] and acidification of the core microenvironment. Typical pH values in these cores range from 6.6 to 7.0 [42-46], which is sufficiently acidic to close most GJs [36-40]. Accordingly, we assume that intercellular communication via GJs is impaired in large tumor spheroids and exclude it from our model. At mesoscales, spheroid cells transmit mechanical stresses through direct contacts, while at macroscales, boundary cells generate interfacial forces that push inward, shaping overall cell morphology and stress distribution. By integrating these multiscale interactions into a unified framework, we can predict the spatial distribution of cell shape and volume, residual solid stress, and intracellular pressure within large tumor spheroids. The governing equations incorporate mechanical force balance at the cellular scale, coupling intracellular pressure to solid and cortical stresses (**Figure 1E**), and include equations describing cellular deformations and displacements (**Figure 1D**), as well as ion and water transport. A detailed description of each component is provided below.

**Figure 1:**
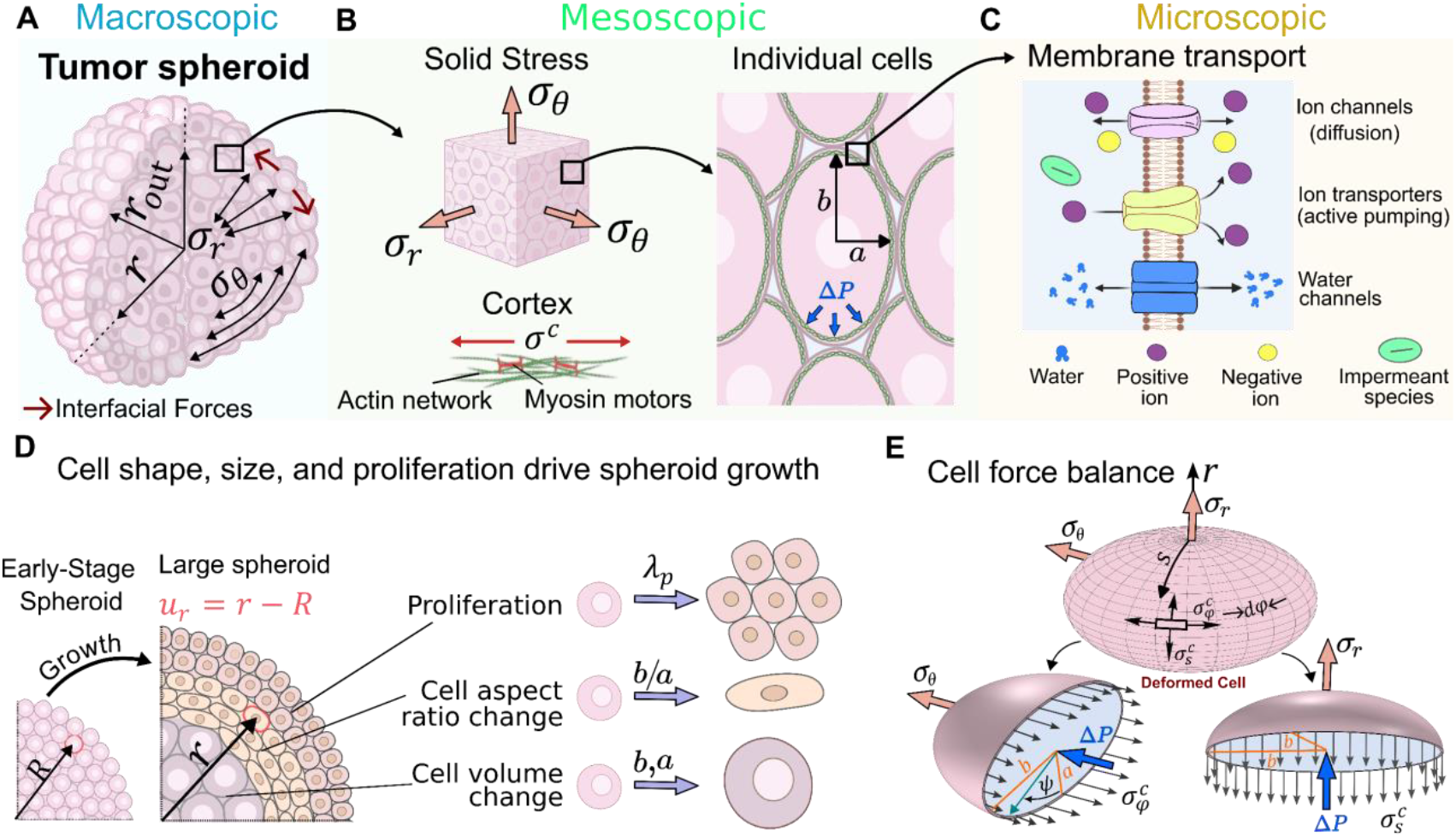
Multiscale mechano-electro-osmotic model linking cellular ion regulation, cortical mechanics, and solid stress generation in tumor spheroids. **A)** Macroscopic: Spheroid geometry in spherical coordinates showing residual solid stresses in the radial (*σ*_*r*_) and tangential (*σ*_*θ*_) directions along with interfacial forces. **B)** Mesoscopic: Transmission of solid stress to cells; the actomyosin cortex generates cortical stress from passive elasticity and myosin activity. Cells are modeled as rotational ellipsoids with radial and tangential semi-axis lengths *a* and *b*, respectively. **C)** Microscopic: Membrane transport regulating cell volume; water moves through aquaporins, ions cross via channels and Na^+^/K^+^ pumps, and impermeant solutes contribute to osmotic pressure. **D)** Early-stage spheroid taken as the reference configuration, with radial displacement defined as *u*_*r*_(*R*) = *r* − *R*. Spheroid growth arises from spatially graded proliferation (*λ*_*p*_) and changes in cell size and aspect ratio (*b*⁄*a*). **E)** Deformed cell subjected to solid stresses, cortical stresses in the meridional 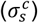 and hoop 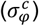 directions, and the hydrostatic pressure difference (Δ*P*) across the membrane.

### 2.1 Solid stress couples to cortical stress and cellular hydrostatic pressure

As cells proliferate or change volume in response to osmotic effects, they exert contact stresses on neighboring cells. To maintain mechanical homeostasis, these external forces must be balanced by internal forces, primarily intracellular hydrostatic pressure and cortical tension. Since the plasma membrane is a fluid-like lipid bilayer that is significantly softer than the underlying actomyosin cortex [47], its contribution to boundary tension is typically negligible.

Under the assumption of circumferential symmetry and employing a spherical coordinate system centered at the spheroid origin (**Figure 1A**), the overall force balance for a representative mesoscale volume element (illustrated as an intermediate scale between the tumor spheroid and individual cells in **Figure 1**) located at radial position *r* can be expressed as:

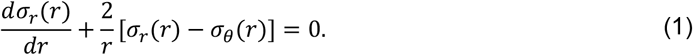

Here, *σ*_*r*_(*r*) and *σ*_*θ*_(*r*) represent the tissue-level solid residual stresses in radial and tangential directions, respectively. These stresses are assumed to uniformly propagate down to the cellular level, influencing cellular shape (**Figure 1E**). As a result, the aspect ratio of individual cells changes through elongation or compression along their primary axes. We model the deformed cell as a rotational ellipsoid with semi-axis lengths *a*(*r*) along the radial direction and *b*(*r*) along the tangential direction of the spheroid (**Figure 1E**). As a result, the condition for force balance on a cell in the radial and tangential directions can be written as (**Figure 1E**):

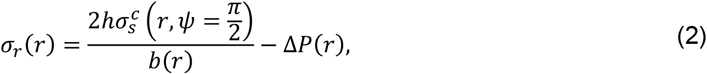

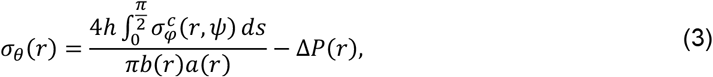

where Δ*P*(*r*) = *P*_*in*_(*r*) − *P*_*out*_(*r*) is the difference in hydrostatic pressure across the cell membrane between the cytosol and the interstitial fluid, and *h* is the thickness of cortex. Additionally, 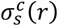 and 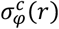 represent the cortical stresses along the cell meridional (*s*) and hoop (*φ*) directions, respectively (See **Figure 1E**). While **Eqs. (2)** and **(3)** describe a force balance at the level of a single cell, they depend on the mesoscopic stress balance (**Eq. (1)**) resulting from the mechanical contact interactions between cells when proliferating within the multicellular environment.

The balance of mechanical forces on the cell, as described by **Eqs. (2)** and **(3)**, indicates that solid stresses are coupled to both the cortical stresses and the hydrostatic pressure difference. The cortical stress components can be expressed as [48]:

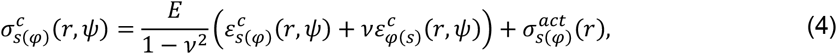

where 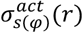 is the active stress generated by myosin activity, and the first term represents the passive stress arising from elastic deformations of the cortex (see **Supplementary Note S1**). Here, *E* and *ν* represent the effective Young’s modulus and Poisson’s ratio of the cortex, respectively, *ψ* denotes the cell polar angle as shown in **Figure 1E**, and 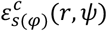 are the cortical strain components. To calculate these strains, we assume that the elongated (ellipsoidal) cells, with semi-axes *a*(*r*) and *b*(*r*), would relax to a spherical shape of radius *a*_0_ under stress-free conditions (see **Figure S1**). Accordingly, the strain components are given by: 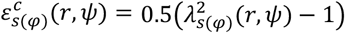, where *λ*_*s*(*φ*)_(*r, ψ*) are the cortical stretches in the cell meridional and hoop directions, respectively, and can be mathematically expressed as (see **Supplementary Note S1** for details):

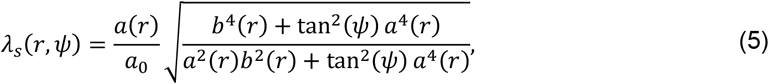

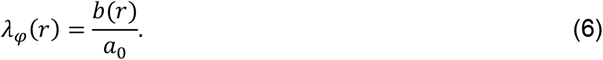

To determine the hydrostatic pressure difference, Δ*P*(*r*), we consider how this pressure gradient is regulated by water exchange across the cell membrane and is therefore coupled to changes in cell volume, *V*(*r*) = 4⁄3 (*πa*(*r*)*b*^2^(*r*)), as discussed next.

### 2.2 Regulation of cell volume through water transport

Water transport across the cell membrane is facilitated by aquaporins, specialized transmembrane proteins that selectively allow the passage of water molecules [31]. This water flux is driven not only by the hydrostatic pressure gradient, Δ*P*(*r*), but also by the transmembrane osmotic pressure difference, ΔΠ(*r*), and can be described by:

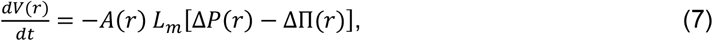

where *A*(*r*) is the cell surface area, and *L*_*m*_ is the membrane’s permeability to water. This relation shows that an elevated hydrostatic pressure inside the cell promotes water efflux, whereas a higher intracellular osmotic pressure favors water influx to dilute the intracellular solute concentration. The osmotic pressure difference can be expressed utilizing the Van’t Hoff equation as: 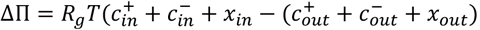, where *c*^+^ and *c*^−^ represent the concentrations of positive and negative ions, *x* denotes the concentration of impermeant solutes, and *R*_*g*_*T* is the product of the ideal gas constant and the absolute temperature. The subscripts “*in*” and “*out*” refer to the intracellular and extracellular compartments, respectively. We study spatial variations in cell volume on the timescale of spheroid growth (days). Since the characteristic timescale of water diffusion across the cell membrane and the associated volume changes (seconds to minutes) is much shorter, cell volume can be considered quasi-static (i.e., *dV*(*r*)⁄*dt* ≈ 0) over the course of spheroid growth. As a result, the hydrostatic and osmotic pressure differences equilibrate, yielding Δ*P*(*r*) = ΔΠ(*r*). Assuming that external solute concentrations remain constant, the transmembrane hydrostatic pressure differences within the spheroid are therefore governed by changes in the intracellular concentrations of ions and impermeant solutes, which are examined next.

### 2.3 Ion concentration in cells: Regulation by ion pumps and channels

Ions can cross the cell membrane through both ion channels and ion pumps [49] (**Figure 1C**). Taking these mechanisms into account, the time rate of change in the number of ionic species can be expressed as [50]:

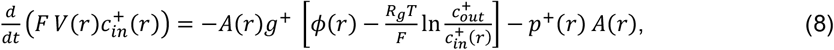

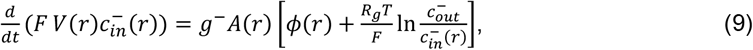

where the first term in each equation describes ion flux through channels, driven by both concentration gradients and the membrane potential *ϕ*(*r*). Here, *g*^+^ and *g*^−^ are the channel conductances for positive and negative ions; and *F* is the Faraday constant. Furthermore, the second term in **Eq. (8)** accounts for active ion transport mediated by Na^+^/K^+^ pumps, which move positively charged ions across the membrane against their electrochemical gradients to maintain cellular ion homeostasis. Here, *p*^+^(*r*) represents the pump rate, defined as the electric current per unit membrane surface area generated by the active efflux of positive charges through ion pumps. Since these pumps are energy-intensive enzymes, consuming approximately 20-30% of a cell’s ATP under basal conditions [51], their activity strongly depends on local oxygen and nutrient availability and therefore varies spatially within large spheroids. This spatial dependence is explored in detail in Section 3.1. Similar to cell volume, changes in the concentration of intracellular ions can also be considered quasi-static (i.e., 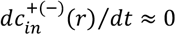) within the spheroid, as their characteristic timescale (seconds to minutes) is considerably shorter than that of spheroid growth (days).

In addition to ions, transmembrane hydrostatic (and osmotic) pressure differences are also influenced by impermeant solutes. These solutes include various molecular species, such as G-actin, organic phosphates, and nucleic acids, which are unable to cross the cell membrane. On average, these species carry negative charges (with valence *z*) and therefore attract counterions to maintain electroneutrality. The electroneutrality condition within the intracellular space can be expressed as:

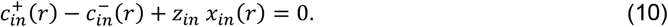

### 2.4 Cell shape, size, and proliferation drive macroscopic deformation in growing spheroids

In the early stages of spheroid formation, growth is primarily driven by uniform cell proliferation [52], as cells throughout the spheroid have ample access to oxygen and nutrients to support division. They typically retain a rounded morphology and maintain near-physiological volume. As the spheroid expands, metabolic gradients gradually emerge, restricting nutrient availability and cell proliferation to the outer layers [52, 53]. Cells located in deeper regions must adapt to this increasingly constrained environment by altering their size, shape, or both. Thus, while early-stage growth is sustained by homogeneous proliferation, growth in larger spheroids with pronounced metabolic gradients involves both spatially heterogeneous proliferation and morphological remodeling. To capture this process, we take the early-stage spheroid as the reference configuration (see **Figure 1D**), where the radial position of a cell is denoted by *R*, and define the radial displacement during growth as *u*_*r*_(*R*) = *r* − *R*. Accordingly, the total deformation in the large spheroid arises from two principal contributions: (1) changes in cell dimensions, encompassing both overall size and aspect ratio, defined as *b*(*r*)⁄*a*(*r*); and (2) displacements of the cells from their reference positions due to cell proliferation. Combining these contributions, the radial and tangential components of the macroscopic stretch arising from the spheroid growth, *λ*_*r*_ and *λ*_*θ*_, can be expressed as (see **Supplementary Note S2** for details):

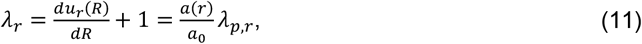

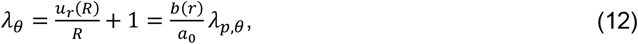

where the first term on the right-hand side of each equation represents the elastic contribution to the total stretch, arising from changes in cell size and shape, while the second term accounts for the proliferation-induced stretch along the radial (*λ*_*p,r*_) and tangential (*λ*_*p,θ*_) directions.

### 2.5 Surface tension determines the level of radial stress at the spheroid boundary

Given that tumor spheroids are microscale soft tissues, interfacial tension plays a key role in their formation, compaction, and, consequently, in the generation of residual solid stresses [54, 55]. This tension primarily arises from the interplay between intercellular adhesion and cortical tension in boundary cells, and it correlates closely with the strength of cell–cell adhesion [56]. Consistent with this, experimental observations [55] show that the surface tension of breast tumor spheroids decreases as cancer cells transition from an epithelial phenotype, characterized by strong cadherin-mediated adhesion, to a mesenchymal phenotype, marked by weaker intercellular contacts. Strong cell–cell adhesions enable boundary cells to redistribute their cortical tension and polarize it tangentially, allowing them to pull on one another along the spheroid surface and generate interfacial tension. Experimental studies in breast [54, 57] and pancreatic [58] cancer spheroids show that this redistribution is particularly pronounced in epithelial aggregates, where boundary cells flatten and elongate, forming a smooth, contractile rim around the spheroid (**Figure 4B**). In contrast, mesenchymal spheroids, which exhibit weaker cell–cell adhesion, are unable to effectively redistribute cortical tension; their boundary cells remain rounded and loosely packed (**Figure 4B**). Since polarization of the cortical contractility is highly confined to the outermost few boundary cell layers [54], we consider its effect through the boundary condition:

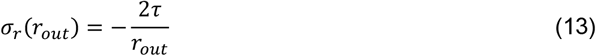

where *τ* is the effective surface tension and *r*_*out*_ is the spheroid radius.

### 2.6 Solving the Mechano-Electro-Osmotic system of equations

In summary, **Eqs. (1)–(3)** and **(7)-(12)** together define a system of nine highly coupled differential-algebraic equations, which serve as the governing equations of the Mechano-Electro-Osmotic model. This system describes nine unknown variables: the solid stress components *σ*_*r*_(*r*) and *σ*_*θ*_(*r*), the cell dimensions *a*(*r*) and *b*(*r*), the transmembrane hydrostatic pressure difference Δ*P*(*r*), the intracellular ion concentrations *c*^+^(*r*) and *c*^−^(*r*), the cell membrane potential *ϕ*(*r*), and the radial displacement of the cell centroid *u*_*r*_(*R*). To eliminate the variable *R* from **Eqs. (11)** and **(12)**, we use the definition *u*_*r*_(*R*) = *r* − *R* to obtain: *du*_*r*_(*R*)⁄*dR* = *du*_*r*_(*r*)⁄*dr* /(1 − *du*_*r*_(*r*)⁄*dr*) and *u*_*r*_(*R*)⁄*R* = *u*_*r*_(*r*)⁄*r* /(1 − *u*_*r*_(*r*)⁄*r*). Furthermore, **Eq. (13)**, together with the condition that the cell centroid displacement remains zero at the spheroid center (*u*_*r*_(0) = 0), provides the boundary conditions for this system. As input values, we specify the spatial distribution of stretches induced by cell proliferation (*λ*_*p,r*_, *λ*_*p,θ*_) and the ion pump rate *p*^+^(*r*), as discussed in the next section. All other model parameters, determined from the literature or our experiments, are summarized in **Table S1** unless otherwise specified. The solutions are computed numerically using Mathematica [59].

## 3. Results

### 3.1 Metabolic dysregulation disrupts ion pump activity and drives core cell swelling

In tumor spheroids, oxygen and essential nutrients like glucose are supplied by the surrounding culture medium and transported inward primarily by diffusion [15]. As outer-layer cells consume these metabolic resources, their availability progressively decreases with depth (**Figure 2A**) due to the combined effects of diffusion and cellular uptake [60, 61]. Experiments [53, 62] show that spheroids larger than ∼200–400 µm in diameter cannot supply enough oxygen through diffusion alone, causing severe hypoxia and metabolic stress in the core. Under these compromised metabolic conditions, mitochondrial ATP production becomes impaired, forcing core cells to rely predominantly on anaerobic glycolysis, a significantly less efficient means of ATP generation. Consequently, ATP levels within the spheroid core decline markedly [63], severely restricting the energy available to sustain Na^+^/K^+^-ATPase activity. In parallel, hypoxia-induced mitochondrial dysfunction elevates the production of reactive oxygen species (ROS) within the core [64]. Increased ROS levels activate signaling pathways known to promote the inactivation, degradation, and endocytosis of Na^+^/K^+^ pumps [65-67], further decreasing their abundance and functionality. Moreover, metabolic dysregulation under hypoxic conditions results in the intracellular accumulation of metabolic byproducts such as lactate [63], lipid droplets [68], and glycogen [53] within the core cells. These metabolic byproducts, mostly carrying negative charges, attract cations into the intracellular space to maintain electroneutrality, further straining the already compromised ion pumps responsible for sustaining physiological ion gradients [69]. Together, ATP depletion, ROS-mediated damage, and accumulation of metabolic byproducts severely disrupt Na^+^/K^+^-ATPase function in core cells (**Figure 2A**). This disruption in ion homeostasis, combined with elevated ROS levels and other metabolic perturbations, such as increased cytosolic calcium, initiates a cascade of detrimental events ultimately driving core cells toward necrosis [70, 71]. Collectively, these biochemical alterations establish a direct link between oxygen availability, ion pump activity, and cell viability within tumor spheroids.

**Figure 2.**
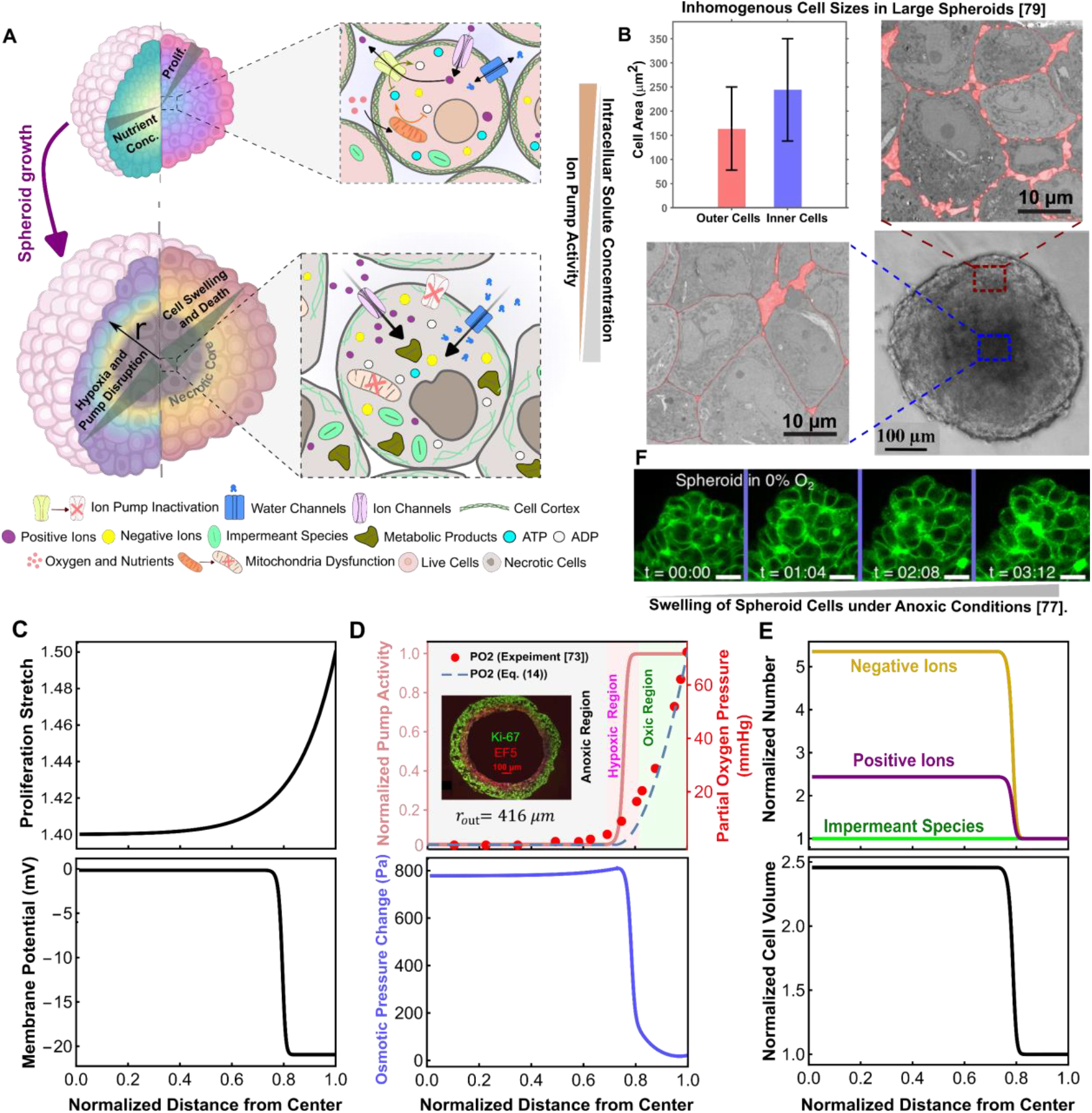
Metabolic gradients and ion pump disruption drive osmotic swelling in tumor spheroid cores. **A)** Schematic depicting the formation of metabolic and oxygen gradients within large tumor spheroids. Reduced nutrient and oxygen availability at the spheroid core impairs mitochondrial ATP production, severely compromising the functionality of Na^+^/K^+^ pumps. This dysfunction leads to ion imbalance, increased intracellular osmotic pressure, and subsequent osmotic swelling of core cells. **B)** High-resolution TEM images of T47D spheroids confirm notable enlargement of necrotic core cells, consistent with predicted osmotic swelling. **C)** Spatial distributions of cell proliferation (upper panel) and model-predicted membrane potential changes (lower panel), illustrating cell membrane depolarization within the spheroid core. **D)** Theoretical and experimental oxygen gradients within spheroids (upper panel), highlighting decreased oxygen partial pressures toward the core that correlate with a sigmoidal reduction in Na^+^/K^+^ pump activity. The corresponding model-predicted intracellular osmotic pressure gradient (lower panel) demonstrates increased pressure within the oxygen-deprived region. *Inset (upper panel):* Immunostaining of DLD1 spheroids [72] reveals that cells are proliferative at the well-oxygenated periphery (Ki-67 positive, green), hypoxic in intermediate layers (EF5 positive, red), and necrotic at the anoxic core. **E)** Predicted accumulation of positive and negative ions (upper panel) and associated cell volume changes (lower panel) across spheroid, illustrating significant ion accumulation and cell swelling confined to the hypoxic central region. **F)** Experimental validation [77] through imaging of small breast tumor spheroids under hypoxic conditions confirms swelling of hypoxic cells (Scale bar = 20 μm).

Experimental measurements and theoretical modeling [72, 73] show that in large spheroids, the radial oxygen gradient is often very steep (**Figure 2D**, upper panel), leading to the formation of an anoxic core. The spatial distribution of oxygen partial pressure, PO_2_(*r*), within the spheroid is given by [72, 74]:

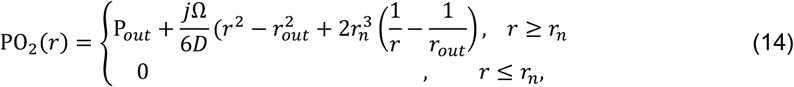

where P_*out*_ is the oxygen partial pressure at the spheroid boundary, *j* is the uniform oxygen consumption rate (OCR), Ω is a constant derived from Henry’s law [72], *D* is the oxygen diffusion coefficient, and *r*_*n*_ denotes the boundary of the anoxic region, defined by the condition PO_2_(*r*_*n*_) = 0 (see [72, 74] for details). Consistent with this oxygen profile, immunostaining of DLD1 (human colorectal carcinoma) spheroids [72] reveals that spheroid cells are proliferative at the well-oxygenated periphery and necrotic at the anoxic core (**Figure 2D**, inset). Motivated by these observations and considering the interplay between oxygenation, cell survival, and Na^+^/K^+^ pump activity, we model the development of the necrotic core by reducing the pump activity (*p*^+^(*r*)) in inner cells as a sigmoid function of oxygen partial pressure:

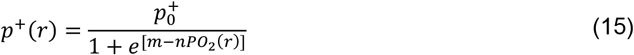

This function (with *m* = 5 and *n* = 1 (mmHg)^−1^ illustrated in **Figure 2D**, upper panel) captures near-physiological pump activity 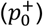 in the outer (proliferative) layers, where the oxygen partial pressure is higher than 10–30 mmHg [72, 75]; complete pump inactivation (*p*^+^ = 0) at the necrotic (anoxic) core, where the oxygen pressure is close to 0 mmHg; and progressively reduced activity in intermediate layers.

Our simulations reveal that with the reduction of Na^+^/K^+^ pump activity in inner cells, the active expulsion of positive (sodium) ions ceases, leading to their intracellular accumulation (**Figure 2E**, upper panel). This buildup of positive ions depolarizes the cell membrane (**Figure 2C**, lower panel), allowing negative (chloride) ions to rush into the cell, as described by **Eq. (9)**. As a result, the total intracellular solute concentration (i.e., 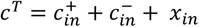) increases in the deprived cells, which, in turn, elevates the intracellular osmotic pressure (Π_*in*_) in these central cells (**Figure 2D**, lower panel). As this osmotic pressure rises within the core, the osmotic gradient across the cell membrane becomes increasingly pronounced, driving water influx from the interstitial space into the cytoplasm to restore equilibrium. This leads to progressive cell swelling in the oxygen-deprived core (**Figures 2A**, and **2E**, lower panel), which in turn elevates intracellular hydrostatic pressure (*P*_*in*_) in this region. In contrast, cells in the outer layers, where pump activity remains largely unchanged, maintain ion homeostasis and osmolarity, preventing swelling and pressure buildup. Consequently, an inhomogeneous cell swelling field emerges across the spheroid (**Figure 2E**, lower panel), with the core exhibiting the highest swelling due to impaired ion regulation and the consequent osmotic dysregulation. These model predictions align closely with multiple independent experimental observations. Microelectrode recordings have revealed depth-dependent depolarization, with membrane potentials approaching zero in the necrotic centers of glioma and hamster tumor spheroids [76]. Hypoxic cell swelling has been documented in small MCF-7 breast spheroids under low oxygen [77]. More importantly, cytoplasmic swelling has also been demonstrated in the central regions of large spheroids derived from human adipose-derived stem cells, with the swollen core expanding markedly as culture oxygenation decreases [78]. High-resolution transmission electron microscopy (TEM) of large T47D tumor spheroids further confirms this phenomenon, revealing substantial enlargement of cells within necrotic cores compared to the outer layers **(Figure 2B**) [79]. Importantly, similar central cell enlargement has also been observed in spheroids derived from lung (SK-MES-1) [80], colon (CT26) [81], and another breast (MDA-MB-436) [82] cancer cell line, demonstrating that core volume expansion is a conserved hallmark of large tumor spheroids, irrespective of tumor type. Collectively, these convergent findings across diverse cancer models provide compelling support for our theoretical results.

Our analysis reveals that, as the spheroid reaches a critical size, rapid depletion of oxygen and nutrients from the spheroid periphery/edge toward the center/core results in a severely deprived core, where metabolic dysregulation impairs ion pump activity, leading to the accumulation of positive and negative ions. This ionic buildup elevates osmotic pressure, driving water influx and resulting in pronounced core cell swelling. Beyond this swelling, the uneven distribution of oxygen and nutrients restricts cell proliferation primarily to the outermost spheroid layers (**Figure 2A**), where metabolic supplies are most accessible, leading to peripheral spheroid growth. To capture this spatially graded proliferation, and guided by experimental measurements [83, 84], we represent the proliferative stretch with an exponential form (**Figure 2C**, upper panel): 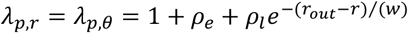. Here, *ρ*_*e*_ reflects the cumulative contribution from the early growth phase, when cell division is nearly uniform across the spheroid. *ρ*_*l*_ denotes the proliferative fraction at the outer boundary in the later phase, when division becomes confined to peripheral cells. The parameter *w* is a characteristic decay length that sets how quickly proliferation decreases from the boundary toward the core as nutrients are depleted (see **Supplementary Note S2** for details). As a consequence of both peripheral growth and core swelling caused by metabolic dysregulation, spatially non-uniform volume changes develop across the spheroid, and mechanical interactions between neighboring cells inevitably arise as cells deform and pack against one another. These cellular-scale forces, in turn, generate and propagate residual solid stresses throughout the spheroid. The resulting stress fields are also influenced by interfacial forces at the spheroid boundary, as previously described (see Section 2.5 and **Eq. (13)**). We next use our model to predict these solid stress fields and cell morphology changes across the spheroid.

### 3.2 Peripheral proliferation cannot explain the emergence of high compressive stresses at the core

Proliferation of cells at the periphery of the spheroid drives local volume expansion. However, this growth is mechanically constrained by neighboring cells and the interfacial forces, generating compressive stresses that propagate inward and shape the overall solid stress distribution. Experimental studies [85-87] have shown that the highest compressive stresses are localized at the spheroid core, with peripheral proliferation thought to be the primary driver of this elevated compression at the center. To systematically evaluate the contribution of proliferation to residual solid stress generation, we first use our mechano-electro-osmotic model to predict the stress profile arising from inhomogeneous volume expansion driven solely by cellular growth. To this end, we eliminate the contribution of osmotic core swelling in our analysis by setting 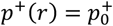 (see **Eq. (15)**), keeping the ion pump activity intact and thereby suppressing water influx within the core.

Our calculations (**Figure 3B**) reveal that constrained peripheral growth generates an inhomogeneous stress distribution, with both tangential and radial compression peaking near the boundary and gradually decreasing toward the center. At the periphery, where cell proliferation is most active, dividing cells exert pressure on one another to accommodate new cells, generating strong compressive forces. However, as we move inward, proliferation becomes increasingly restricted, leading to a reduction in growth-driven expansion and a weakening of compressive forces between cells. Consequently, the solid stresses decline with depth and eventually stabilize in the core, where cell division ceases. Notably, this predicted stress distribution stands in stark contrast to experimental observations [81, 85-87], which consistently show that solid compressive stresses intensify toward the core rather than diminishing.

**Figure 3:**
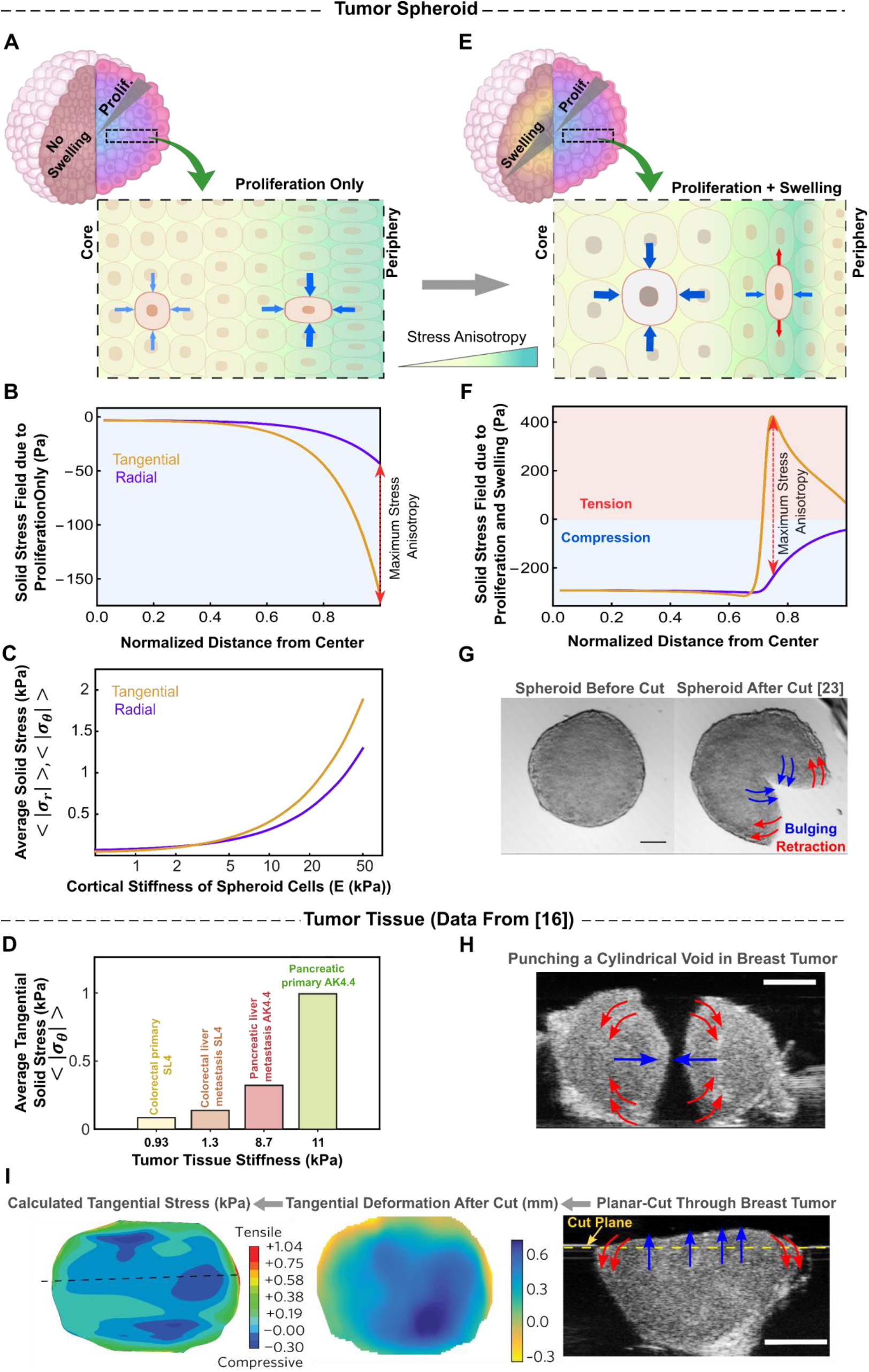
Osmotic core swelling, rather than peripheral proliferation, drives the emergence of residual solid stresses in tumor spheroids. **A, E)** Schematics illustrating how the residual solid stress profile differs when only proliferation-driven volume expansion is considered (left), versus when osmotic swelling due to pump failure is also included (right). **B)** Model predictions with peripheral proliferation alone show compressive stresses peaking at the periphery and decreasing toward the core, in contrast to experimental observations. **C)** Predicted mean solid stress magnitude increases with spheroid cell cortical stiffness. **D)** Cutting-based stress-release measurements in excised mouse tumors (data from [16]) show that the average tangential solid stress is on the order of 0.1–1 kPa and tends to increase with tissue stiffness across tumor types indicated. **F)** Incorporating osmotic swelling into the model shifts the peak compressive stress to the core and generates peripheral tangential tension, consistent with measurements in multicellular spheroids. **G)** Partial cuts through HCT116 spheroids [23] confirm this prediction, exhibiting central bulging (indicative of compression) and peripheral retraction (indicative of tension). Scale bar: 100 μm. **H**) Needle-biopsy stress-release assay in an MMTV-M3C breast tumor ([16]) shows narrowing under compression and widening under tension. Scale bar, 2 mm. **I**) Planar-cut assay in the same tumor model ([16]) shows bulging at the center and retraction at the periphery; the measured deformation field and inferred tangential stress map are shown. Scale bar, 2 mm.

**Figure 4:**
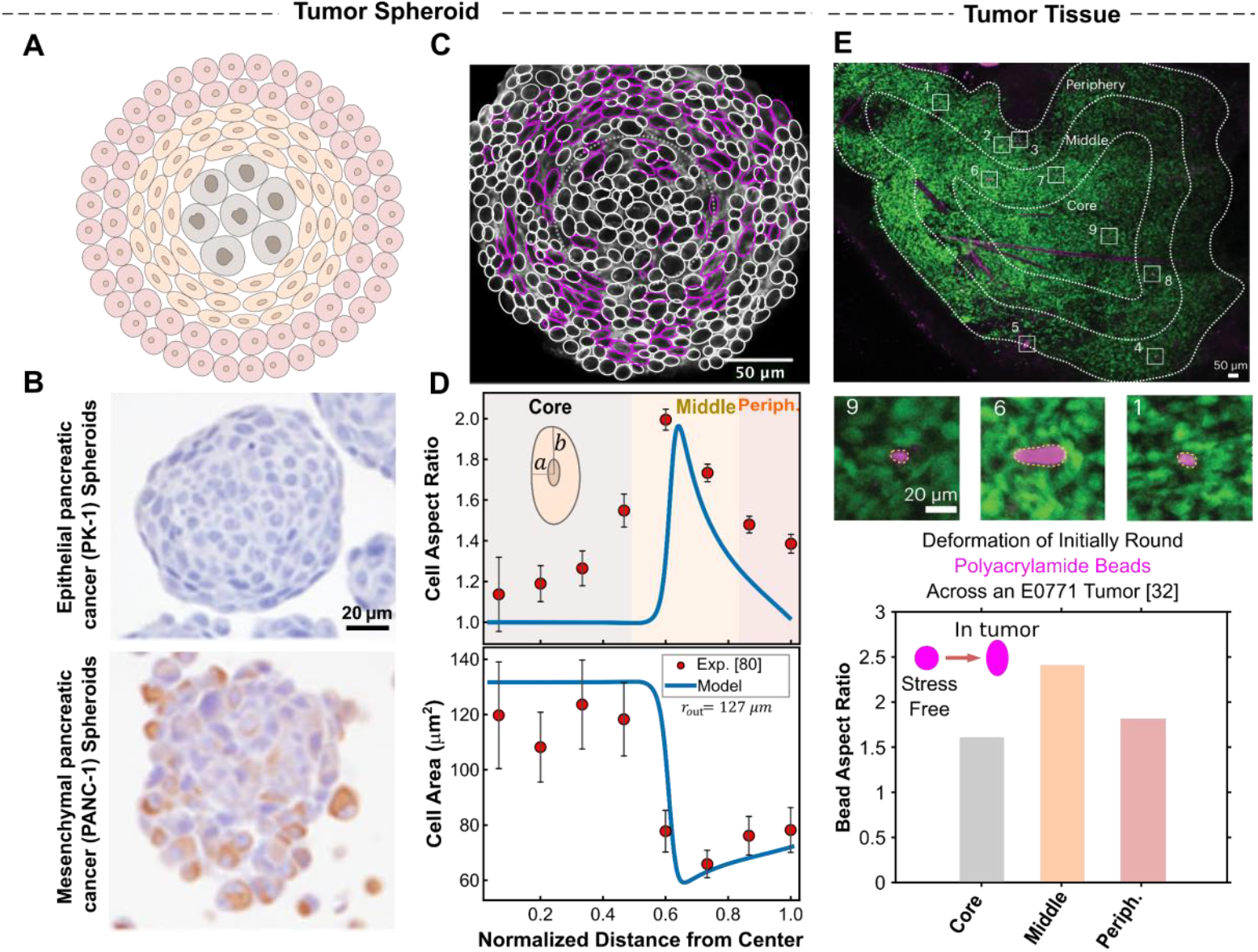
Model predicts distinctive cell shape changes primarily driven by osmotic swelling in tumor spheroids. **A)** Spatial organization of tumor spheroids into three morphologically distinct zones: an isotropically swollen core, a highly elongated intermediate region, and a slightly elongated proliferative periphery. **B)** Boundary cell shape is governed by interfacial forces, remaining rounded in mesenchymal spheroids with weak surface tension but becoming tangentially elongated in epithelial spheroids with strong surface tension, as observed in pancreatic small cancer spheroids [58]. **C)** Equatorial-plane image of a CT26 (mesenchymal) spheroid [81] showing pronounced cell elongation localized primarily in the intermediate zone, consistent with theoretical predictions. **D)** Quantitative comparison between experimental measurements (mean ± s.d.) and model predictions of cell shape (top) and area (bottom) variations in CT26 tumor spheroids, demonstrating strong agreement and supporting the model’s accuracy. **E)** In vivo evidence from E0771 breast tumors [32] showing polyacrylamide beads embedded in the tissue becoming elongated predominantly in intermediate regions, supporting the model’s ability to capture key biophysical features of tumor tissue.

To reconcile this discrepancy, Dolega et al. [81] proposed that cell stiffness is anisotropic within the spheroid and drives the accumulation of proliferation-induced solid stress at the core. They showed that when spheroid cells are stiffer in the radial direction than in the ortho-radial plane, solid stress components scale with depth as a power law, *σ*_*r*(*θ*)_ ∝ *r*^−*β*^, *β* > 0. While conceptually intriguing, this assumption of stiffness anisotropy leads to physically inconsistent predictions (see **Supplementary Note S3** for details). Specifically, it results in a singular (unbounded) solid stress field at the spheroid center, which is physically unrealistic. More importantly, it suggests that stress anisotropy (*σ*_*r*_ − *σ*_*θ*_) increases toward the core, contradicting recent experimental findings [85, 86], which reveal that stress within the core is nearly isotropic. Collectively, these inconsistencies indicate that peripheral growth alone cannot account for the large compressive stresses observed at the spheroid core. Rather, this underscores the necessity of additional mechanisms that shape the residual solid stress distribution within tumor spheroids.

### 3.3 Osmotic cell swelling is the primary driver of elevated residual solid stress in tumor spheroids

As outlined in Section 3.1, metabolic dysregulation in large spheroids not only drives peripheral growth but also disrupts ion pump activity within the core. This disruption elevates osmotic pressure and leads to cell swelling in the central region (**Figure 2**). Because the expansion of core cells is mechanically constrained by the surrounding viable layers, it introduces an additional source of compressive stress within the spheroid. Surprisingly, despite their potential significance, the effects of swelling-induced forces on the spheroid’s solid stress fields have not been previously investigated. To evaluate their impact, we incorporate core swelling into our simulations, alongside peripheral proliferation, by enabling ion pump disruption in the core (see **Eq. (15)**).

We find that including core swelling leads to a substantial shift in stress distribution: the highest compressive stresses, both tangential and radial, relocate from the periphery to the spheroid center (**Figure 3F**, compared to **Figure 3B**). This redistribution arises from large compressive forces generated by the confined swelling of the core. As the core expands, it exerts contact stresses on the surrounding viable cell layers, generating radial compressive forces at their interface that propagate throughout the spheroid. Within the core, these forces give rise to a large, homogeneous hydrostatic stress field due to its spherical symmetry and uniform disruption of ion pump activity. Outside the core, however, radial compression gradually decays. This decay occurs because the compressive forces stretch successive cell layers tangentially, inducing tension in that direction (**Figure 3F**). The resulting tangential tension counteracts radial compression, progressively weakening it with distance from the core. Consequently, far from the core, the stress field due to osmotic swelling weakens, while compressive stress from peripheral proliferation becomes dominant. Together, these findings show that osmotic swelling induces high compressive stresses in the spheroid core, balanced by tangential tension in the outer layers (**Figure 3F**). Notably, this predicted stress profile agrees well with the experimental measurements of residual solid stress in tumor spheroids derived from HCT116 colon carcinoma cells [23, 24]. In these experiments, partial cuts through the spheroid center led to noticeable bulging at the core and retraction at the boundary (**Figure 3G**), revealing tangential stresses that are compressive in the central region and tensile in the periphery.

Interestingly, a similar compressive–tensile solid-stress profile has been observed ex vivo using cutting-based stress-release assays applied to primary and metastatic breast tumors, as well as multiple other tumor tissues [16]. **Figures 3H** and **3I** show residual solid-stress-induced deformation in an MMTV-M3C primary tumor following needle biopsy and planar cutting. In the needle-biopsy assay, residual stress is relieved by punching a cylindrical void, and the resulting diameter changes report the local stress state, with narrowing under compression and widening under tension (**Figure 3H**). In the planar-cut assay, releasing mechanical confinement along a defined cut plane produces a characteristic deformation pattern, with bulging at the tumor center and retraction at the periphery, consistent with central compression and peripheral tangential tension (**Figure 3I**). Across these samples and the additional tumor tissues summarized in **Figure 3D**, these cutting-based measurements indicate that the average tangential solid stress (< |*σ*_*θ*_| >; **Supplementary Note S5**) in mouse tumors is on the order of 0.1–1 kPa and increases with stiffness (**Figure 3D**). Notably, our model predicts the same stress scale and captures a similar increase with spheroid cell stiffness (**Figure 3C**), reinforcing that tumor spheroids can reproduce solid-stress levels measured in tumor tissues, consistent with recent intravital measurements [32].

Overall, our analysis reveals that residual solid stress in tumor spheroids primarily arises from two distinct local volume expansion mechanisms: peripheral proliferation and core swelling. Proliferation-driven stress results from mechanical forces generated by dividing cells at the periphery, whereas swelling-driven stress stems from the constrained expansion of the core. Notably, proliferation alone cannot reproduce the experimentally observed stress distribution. Instead, core swelling emerges as the dominant driver of elevated compressive stress at the center and tensile tangential stress at the periphery. In the following section, we examine how these stresses influence cell morphology by analyzing the contributions of proliferation- and swelling-dominated regions.

### 3.4 Core swelling regulates the spatial organization of cell shape across spheroids

Our results for the solid stress field (**Figures 3B** and **3F**) indicate that incorporating osmotic cell swelling not only shifts the location of maximum compressive stresses toward the spheroid center but also markedly alters the anisotropy of the stress field (i.e., |*σ*_*θ*_ − *σ*_*r*_|). While proliferation-driven forces localize peak stress anisotropy near the spheroid boundary (**Figure 3B**), the addition of swelling-induced forces shifts the maximum anisotropy toward the core (**Figure 3F**). Notably, our theoretical analysis (**Supplementary Note S4**) reveals a direct relationship between local stress anisotropy and cell shape changes in large spheroids, expressed as deviations of the aspect ratio from unity: (*σ*_*θ*_ − *σ*_*r*_) ∝ (*b*/*a* − 1). This correlation suggests that regions experiencing the highest stress anisotropy also undergo the most pronounced cell elongation. In this context, cell shape becomes more than a morphological descriptor; it serves as a simple yet effective spatial readout of the underlying stress field inside the spheroid. Importantly, experimental measurements in CT26 spheroids [81] reveal that the most pronounced cell elongation occurs in intermediate regions, well away from the spheroid boundary, and predominantly in the tangential direction (**Figures 4C** and **D**). This observation further challenges the notion that peripheral proliferation alone shapes the stress landscape, as such a mechanism would instead predict maximal elongation at the boundary and in the radial direction (**Figures 3A** and **B**).

To gain further insight into the spatial patterning of cell morphology, we then employ our model to examine how core swelling drives regional variations in cell shape across the spheroid. In the early stages of growth, spheroid cells have ample access to oxygen and nutrients, enabling them to maintain ionic homeostasis. At this stage, solid stress remains minimal due to low cell density and limited mechanical confinement. As the spheroid enlarges, however, metabolic gradients emerge, causing cellular swelling in the core while restricting proliferation to the periphery. The resulting gradients in cell size and number compel spheroid cells to remodel their morphology accordingly (see **Eqs. (11)** and **(12)**). Our model predicts that, in large spheroids, these constraints give rise to three distinct morphological zones (**Figure 4A**), each characterized by unique patterns of cell shape.

#### 1) Swelling dominant core

Within the spheroid core, where cell swelling is maximal, our model predicts that the swelling remains isotropic and spatially uniform (**Figure 4D**, lower panel), owing to the region’s spherical symmetry and evenly distributed osmotic pressure (**Figure 2D**, lower panel). As a result, the solid stress in the core is also isotropic (*σ*_*θ*_ = *σ*_*r*_), allowing cells in this zone to expand while maintaining a rounded morphology (*b*/*a* = 1) (**Figure 4D**, upper panel).

#### 2) Proliferation dominant periphery

At the periphery, where spheroid growth is primarily accommodated by cell proliferation, cells are well-nourished and thus maintain their physiological volume. Our results show that stress anisotropy in this region gradually decreases toward the spheroid boundary. Consequently, cells near the boundary are slightly elongated (*b*⁄*a* ∼1) along the direction of lower compression (**Figure 4D**, upper panel). This predicted low stress anisotropy and slight cell elongation hold true at the boundary of mesenchymal spheroids (e.g., CT26 spheroids [88] in **Figure 4D**), where intercellular adhesions are weak, resulting in minimal changes to the shape of boundary cells. In epithelial spheroids, however, strong E-cadherin-mediated adhesions enable boundary cells to polarize their cortical tension along the spheroid boundary, generating significant interfacial tension that flattens and elongates them tangentially (**Figure 4B**).

#### 3) Intermediate region

Located between the swelling core and the proliferative periphery, cells in this zone experience neither sufficient osmotic pressure to drive maximal swelling nor adequate nutrient supply to sustain active proliferation. Instead, they undergo the most pronounced shape changes as a consequence of spheroid growth at the periphery and cell expansion at the core. As a result, these cells become markedly elongated in the tangential direction and compressed radially, exhibiting significantly high aspect ratios (*b*⁄*a*) (**Figure 4D**, upper panel), thereby facilitating radial expansion of intermediate layers. This shape polarization arises primarily from radial pressure exerted by the swelling core on the inner side and the proliferating peripheral cells on the outer side, effectively “sandwiching” the intermediate region. These radial pressures generate a highly anisotropic field of residual solid stress in the intermediate region (**Figure 3E**) with higher compression in the radial direction (*σ*_*θ*_ > *σ*_*r*_). This anisotropy peaks near the core boundary and gradually diminishes toward the periphery, leading to maximal tangential elongation just outside the core, which then gradually decreases toward the periphery (**Figure 4D**, upper panel).

Together, our model predicts a distinctive pattern of cell shape variation across the spheroid in the presence of core swelling: cells remain nearly round at the core, exhibit pronounced tangential elongation in intermediate regions and show reduced elongation toward the periphery (**Figure 4A**). This predicted spatial organization of cell shapes closely matches experimental observations in CT26 spheroids [81]. To quantitatively validate our model, we analyzed an equatorial-plane image of the CT26 spheroid from [81], extracting spatial variations in cell size and shape across the aggregate (see *Methods* and the following section for image analysis details). Notably, the model reproduces the spatial gradients in both cell size and shape observed in this mesenchymal spheroid (**Figure 4D**), using physiologically relevant model parameters (see **Table S1** and **Supplementary Note S5**). These results support the validity of our model and underscore the critical role of osmotic cell swelling in shaping the spheroid’s solid stress landscape. Interestingly, a similar biphasic pattern of solid stress anisotropy has also been observed *in vivo* in mesenchymal-like E0771 breast tumors [32, 89], inferred from measured aspect ratios (**Figure 4E**, lower panel) of (initially spherical) polyacrylamide beads embedded within the tumor tissue (**Figure 4E**, upper panel). This finding further reinforces that our tumor spheroid model faithfully captures key aspects of tumor biophysics.

### 3.5 Experiments on triple-negative breast cancer cells (MDA-MB-231) validate model-predicted cell morphology changes

To further test our model’s predictions of cell morphology changes, we next generated spheroids from MDA-MB-231 breast cancer cells, which exhibit a mesenchymal phenotype [90], by seeding cells in ultra-low-attachment round-bottom plates under stress-free culture conditions. Cells formed compact spheroids within ∼3 days and continued to grow through days 11 and 14, reaching an average radius of ∼200 μm, with cells present in both the core and periphery. These spheroids were then fixed, cleared, and imaged (see *Methods*). We stained filamentous actin using phalloidin (shown in grayscale; **Figures 5A, C, S6A**, and **S7**), and assessed cell proliferation using Ki-67 staining (cyan; **Figure S6C**). Our analysis revealed that cell proliferation is predominantly restricted to the outer layers of the spheroid, consistent with our model assumptions and in agreement with previous findings [53, 91]. We next analyzed the size (area) and shape (aspect ratio) of cells within the spheroids using AI-based cell segmentation on confocal microscopy images of middle-plane cross-sections (**Figures 5A, C, S6B** and **S7**). Specifically, we leveraged the recently developed CellSAM model [92] which integrates both the nuclear (Sytox) and cell-boundary (phalloidin) channels to accurately segment individual cells. These segmentations enabled quantitative analysis of the spatial distributions of cell area and aspect ratio across the spheroid (see *Methods* for details). We assumed that cell organization within the spheroids exhibits spherical symmetry; that is, cell deformations along the polar and azimuthal directions are equivalent. This assumption allowed us to use cell cross-sectional area as a proxy for the cell volume. We further estimated the cell aspect ratio (i.e., *b*⁄*a*) as the ratio of the cell axis in the tangential plane (2*b*) to that in the radial plane (2*a*), as illustrated in **Figure 4D**

We found that in spheroids grown until day 11, both cell volume and shape strongly correlated with radial position within the cluster (**Figure 5B**). Cells in the core (approximately the inner 35% of the spheroid radius) were nearly uniform in size, with a cross-sectional area of ∼380 μm^2^, significantly larger than those in the periphery (outer 30% of the spheroid radius), where the average cross-sectional area was ∼260 μm^2^. Additionally, while cells in the core and near the boundary remained round (aspect ratio ∼1), those in the intermediate layers became markedly compressed in the radial direction and elongated tangentially, reaching aspect ratios as high as 1.5–2 (**Figure 5B**), as highlighted by magenta boundaries in **Figure 5A**. Interestingly, these observations of cell size and shape are in strong quantitative agreement with our model predictions (**Figure 5B**), obtained using physiologically grounded parameters (see **Table S1** and **Supplementary Note S5**). This concordance supports the validity of our model.

**Figure 5:**
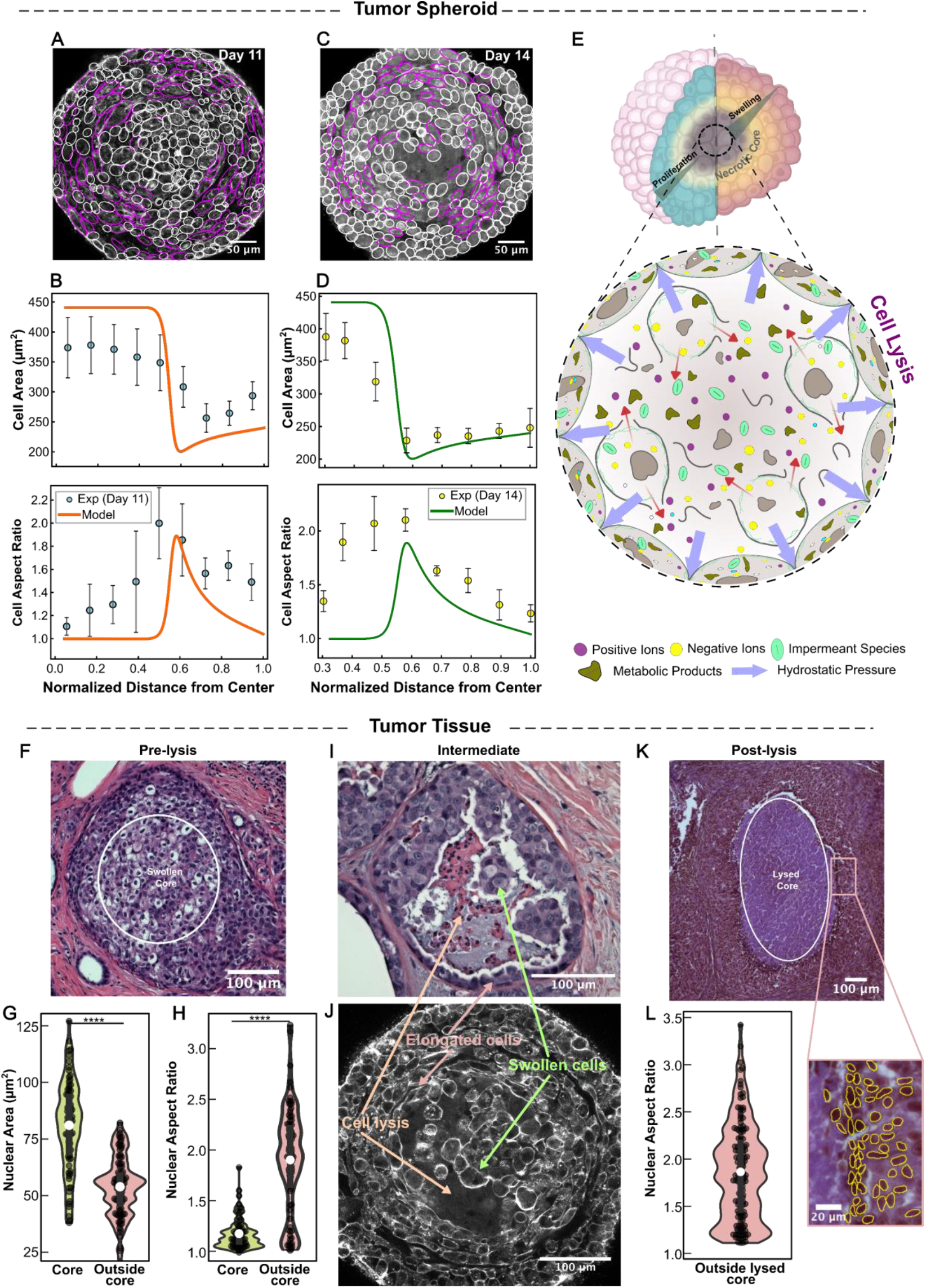
Experimental validation of model-predicted cell morphology changes in large tumor spheroids using triple-negative breast cancer cells (MDA-MB-231) and patient-derived DCIS lesions. **A)** Representative confocal mid-plane image of an MDA-MB-231 spheroid cultured for 11 days and stained for F-actin (phalloidin, shown in grayscale). Magenta outlines denote cells with high aspect ratio (AR > 1.7), highlighting tangentially elongated cells in the intermediate region of the spheroid. **B)** Quantitative spatial analysis of cell cross-sectional area (top) and aspect ratio (bottom) for day-11 spheroids reveals isotropic, homogeneous swelling at the spheroid core, maximal tangential elongation in the intermediate zone, and reduced elongation near the periphery, matching model predictions. **C)** Confocal image of a 14-day-old spheroid showing extensive lysis of necrotic cells at the core, evidenced by large cell-free regions. Magenta outlines again denote cells with high aspect ratio (AR > 1.7), highlighting tangentially elongated cells. **D)** Model-predicted spatial profiles of cell morphology following lysis of necrotic core cells remain consistent with experimental observations, showing that the release of intracellular contents after lysis of necrotic cells sustains osmotic swelling and maintains compressive stress at the core. **E)** Schematic illustrating how necrotic cores transform into polyelectrolyte-rich environments following cell lysis, resulting in persistent osmotic pressure at the spheroid center. **F)** Representative H&E-stained section of a DCIS lesion showing a central region of enlarged, rounded cells forming a swollen core (outlined). **G)** Quantification of nuclear cross-sectional area demonstrates increased nuclear (cell) size in the core relative to the surrounding region. Each dot represents a segmented nucleus, *n = 72* (core) and 90 (outside core). **H)** Nuclear aspect ratio quantification shows that nuclei are relatively round within the swollen core but become highly elongated in the surrounding region. Each dot represents a segmented nucleus, *n = 72* (core) and 90 (outside core). **I, J)** Comparison of intermediate-stage morphology in DCIS tissue sections and in vitro tumor spheroids. **I)** H&E-stained DCIS section showing cell swelling and localized cell lysis in the duct center, surrounded by tangentially elongated viable cells. **J)** In vitro tumor spheroid at a comparable stage, exhibiting similar spatial organization, with central cell swelling and lysis accompanied by surrounding tangential elongation, highlighting similar underlying processes across the patient-derived tissue and tumor spheroids. **K)** DCIS lesion exhibiting a lysed core (outlined) surrounded by compressed viable epithelial layers; inset shows tangentially elongated nuclei adjacent to the necrotic lumen. **L)** In lesions with lysed cores, nuclei in the surrounding viable rim exhibit elevated aspect ratios, consistent with strong radial compression exerted by the swollen core. Each dot represents a segmented nucleus, *n = 141* (outside lysed core). Quantification in panels **B** and **D** (mean ± s.d.) is based on *n* = 3 independent spheroids per time point. DCIS analyses are based on *n* = 3 independent lesions per condition (lesions with core swelling for F–H; lesions with prominent lysis for K–L). ^****^ indicates p < 0.0001. Additional representative images with magnified sections for each case are provided in the **Supplementary Information (Figures S6, S7 and S11)**.

Importantly, when we allowed the spheroids to grow in media for 14 days, immunostaining and image analyses (**Figure 5C**) revealed that core cells lost membrane integrity, as indicated by cell-free areas in this region. These findings suggest that excessive water influx due to metabolic dysregulation leads to widespread lysis of necrotic core cells, causing the uncontrolled release of their contents, particularly charged species, into the surrounding extracellular milieu (**Figure 5E**). This conclusion aligns with experimental findings in human and mouse tumors [93], which show that necrosis in these cell aggregates releases intracellular potassium ions into the extracellular fluid. The breakdown of the plasma membrane in the central necrotic cells causes their cytoplasmic constituents to mix with the surrounding media, transforming the initially highly heterogeneous multicellular structure of the region into a more homogeneous, gel-like, multi-phase microenvironment [94], characterized by central cell-free regions in our images (**Figure 5C**). We next extend our theoretical framework to study how these compositional changes in necrotic cores impact residual solid stress fields and cell shape changes within tumor spheroids.

### 3.6 Model-predicted solid stress and cell morphology patterns persist after lysis of necrotic cells

We model the necrotic core after the lysis of its constituent cells as a polyelectrolyte solution primarily composed of water, mobile ions, and, most importantly, negatively charged macromolecules derived from the released cellular contents (see **Supplementary Note S6** for details). Upon cell lysis, a broad spectrum of macromolecular debris is released, many of which are negatively charged [95, 96]. These macromolecules span sizes from nanometers to microns, while pore sizes in the spheroid core are typically limited to 15–50 nm under physiological conditions [97-99]. Consequently, large debris components, such as sub-micron DNA fragments [100], cannot freely diffuse through the narrow interstitial spaces between adjacent cells and thus become trapped within the core (**Figure 5E**). To neutralize the negative charges fixed to these macromolecules, positive ions also remain within the core, resulting in a higher ion concentration in this central region compared to the surrounding microenvironment. This difference in ion concentration gives rise to the well-known Donnan osmotic pressure [101], which prevents water from escaping the core after lysis, keeping it in a swollen state. Additionally, based on Ki-67 staining at earlier time points in our experiments (**Figure S6**), we assume that cell proliferation remains predominantly localized to the spheroid periphery following lysis of core cells. Therefore, the driving forces for solid residual stress generation and cell shape changes within the spheroid, namely, core swelling and peripheral growth, remain essentially unchanged before and after lysis of necrotic cells. Prior to lysis, individual cells in the deprived core swell and push against the surrounding cell layers. However, post-lysis, the elevated Donnan osmotic pressure retains water within the core, maintaining its enlarged volume and creating similar central compressive forces (**Figure 5E**).

Consequently, our extended model predicts solid stress fields (**Figure S8**) and cell morphology changes (**Figure 5D**) that are qualitatively similar to our earlier findings for spheroids without lysis (see Sections 3.3 and 3.4). The stress field within the core remains highly compressive, uniform, and isotropic due to confined swelling, spherical symmetry, and even distribution of trapped charges following widespread lysis. Outside the core, however, these compressive stresses become anisotropic (**Figure S8**), with maximum compression primarily in the radial direction across the viable cell layers. This higher compression in the radial direction flattens viable cells radially and elongates them tangentially (**Figure 5D**). Cell tangential elongation is maximum in intermediate regions and decays with radial distance from the center. Notably, these model predictions for cell elongation accurately follow our experimental measurements in 14-day spheroids (with widespread lysis) when physiologically relevant model parameters are chosen (see **Supplementary Notes S5** and **S6** for details), further confirming the applicability of our theoretical framework.

Together, these results demonstrate that our model systematically integrates biological and mechanical processes within the spheroid, revealing that osmotic swelling is a primary driver of solid stress accumulation in the spheroid core, even after cell lysis. This mechanistic insight bridges the gap between cellular-scale metabolic events and tissue-scale mechanical phenomena, providing a comprehensive framework for understanding stress evolution at different stages of tumor spheroid development.

### 3.7 Model-predicted spatial organization of cell morphology is recapitulated in patient-derived DCIS tissue

To provide an independent validation in primary human tissue, we analyzed patient-derived human ductal carcinoma in situ (DCIS) lesions to test the model’s predicted spatial organization of cell morphology in a native tumor microenvironment. DCIS is a particularly suitable system because neoplastic epithelial cells proliferate within the duct, while the vasculature remains confined to the surrounding stroma. This geometry naturally creates diffusion-limited gradients of oxygen and nutrients across the intraductal tumor mass [102]. Intraductal growth can produce epithelial thicknesses exceeding ∼0.125–0.160 mm, which surpass the typical oxygen diffusion distance in tissue. As a result, a well-oxygenated rim forms near the basement membrane, whereas conditions become progressively hypoxic toward the duct lumen [103]. Under these conditions, hypoxia promotes a metabolic shift toward glycolysis, leading to increased lactic acid production and progressive acidification toward the duct center [102, 103]. Together, the resulting radial gradients in oxygen, nutrients, and pH closely resemble the diffusion-limited microenvironment that develops in large tumor spheroids. Consistent with this diffusion-limited organization, representative DCIS lesions show enlarged, rounded cells and localized lysis near the duct center, surrounded by tangentially elongated cells (**Figure 5I**). This spatial organization closely resembles that observed in our in vitro spheroids at comparable stages (**Figure 5J**), suggesting a conserved, ordered pattern of central swelling and lysis accompanied by surrounding tangential elongation across the two systems.

To quantify these patterns in patient-derived clinical specimens, formalin-fixed paraffin-embedded DCIS specimens were sectioned, H&E stained, and imaged at high resolution. We used a combination of the StarDist algorithm [104] and manual segmentation in Fiji to generate spatially resolved statistics of nuclear size and shape across the tumor epithelium. Because cell boundaries are often indistinct in H&E sections, nuclear cross-sectional area and nuclear aspect ratio were used as proxies for cellular size and shape, consistent with established quantitative histopathology approaches [105]. Two representative lesion states were analyzed: lesions without histologically visible lysis (**Figure 5F**) and lesions with prominent central lysis (**Figure 5K**). This allowed us to assess cell morphology gradients in both a pre-necrotic diffusion-limited context and in an advanced necrotic lesion.

In lesions without lysis, the nuclear area increased toward the duct interior (**Figure 5G**), indicating larger cells in the hypoxic core than in the well-oxygenated periphery. This radial increase in nuclear (cell) size is consistent with our spheroid measurements (**Figure 5B**) and with model predictions that diffusion-limited metabolic compromise and ionic dysregulation promote central cell swelling. Nuclear shape showed a complementary spatial pattern: nuclei were comparatively round within the swollen core but became highly elongated in the surrounding region (**Figure 5H**). This peripheral elongation is consistent with a model-predicted zone of elevated stress anisotropy, which drives tangential cell elongation around the core. In lesions with a lysed core, nuclei in the surrounding viable cells were strongly elongated and tangentially aligned (**Figure 5L**), again consistent with our in vitro observations (and model predictions; **Figure 5D**). This tangential elongation implies a highly anisotropic stress field at the core boundary, with higher compressive stress in the radial direction. Within our framework, such radial compression arises from an outward mechanical load generated by Donnan-type osmotic pressure in the swollen, osmotically imbalanced necrotic core (**Figure 5E**), which continues to exert radial compressive stresses on the surrounding viable tissue.

Together, the spatial distributions of nuclear (cell) size and shape in DCIS lesions (**Figures 5F–L, Figure S11**), combined with independent evidence for diffusion-limited hypoxia, glycolysis-driven acidification, and reduced ATP availability toward the duct interior, provide validation for key predictions of our mechano–electro–osmotic framework in a native tissue context. This convergence of theory, in vitro spheroid experiments, and patient-derived DCIS histopathology supports the relevance of our model’s mechanistic assumptions and demonstrates that the predicted cell morphology patterns can also arise in clinically relevant tumor microenvironments.

## 4. Discussion

In this study, we developed a mechano-electro-osmotic (MEO) model (**Figure 1**) to elucidate how residual solid or contact stresses between cells are generated and transmitted within large (diameter > 200-400 *μm*) tumor spheroids with pronounced metabolic gradients. Using a bottom-up approach that enforces mechanical and electrochemical balances, we demonstrated that these contact stresses are regulated by multiscale processes: at the cellular level, by actomyosin contractility and volume regulation via fluid and ion transport; and at the tissue level, by cell proliferation and surface tension. Importantly, these cellular processes are energetically demanding and therefore tightly modulated by the local availability of oxygen and nutrients, which sharply decline from the spheroid periphery to the core due to limited diffusion and sustained consumption. Under basal conditions, protein synthesis accounts for the largest share of ATP consumption, followed by Na^+^/K^+^-ATPase activity, with smaller contributions from actomyosin contractility and other metabolic processes [51]. Thus, in the hypoxic core, where cell cycle progression and protein synthesis are suppressed, ion pumps constitute the dominant residual ATP demand and become the most vulnerable target of metabolic stress. Accordingly, we showed that metabolic dysregulation within the spheroid core not only restricts cell proliferation to peripheral regions but also impairs ion pump activity, leading to intracellular ion accumulation, elevated osmotic pressure, water influx, and marked cellular swelling in this region (**Figures 2, 6C**, and **6D**). By contrast, cells in the outer layers maintain ion homeostasis and osmotic balance due to preserved pump activity, thereby preventing swelling and pressure buildup. Consequently, an inhomogeneous swelling field develops across the spheroid, with the core exhibiting the most pronounced swelling. This model prediction aligns with our experimental findings (**Figure 5A-D**) and with previous observations in tumor spheroids derived from lung [80], colon [81], and breast [79, 82] cancer cells, all of which demonstrate that core cells are markedly larger than those in the outer layers, suggesting that this phenomenon may be independent of tumor type. Consistent with these trends, our analysis of human DCIS tissue likewise revealed a comparable radial organization of cellular size, reinforcing that the mechano-electro-osmotic mechanisms identified in spheroids are also applicable in the native tumor epithelium (**Figure 5F-L**). To accommodate both peripheral growth and core swelling driven by metabolic dysregulation, cells within the spheroid must exert forces against one another, adjusting their volumes and shapes, thereby generating and propagating residual solid stresses throughout the tissue.

**Figure 6:**
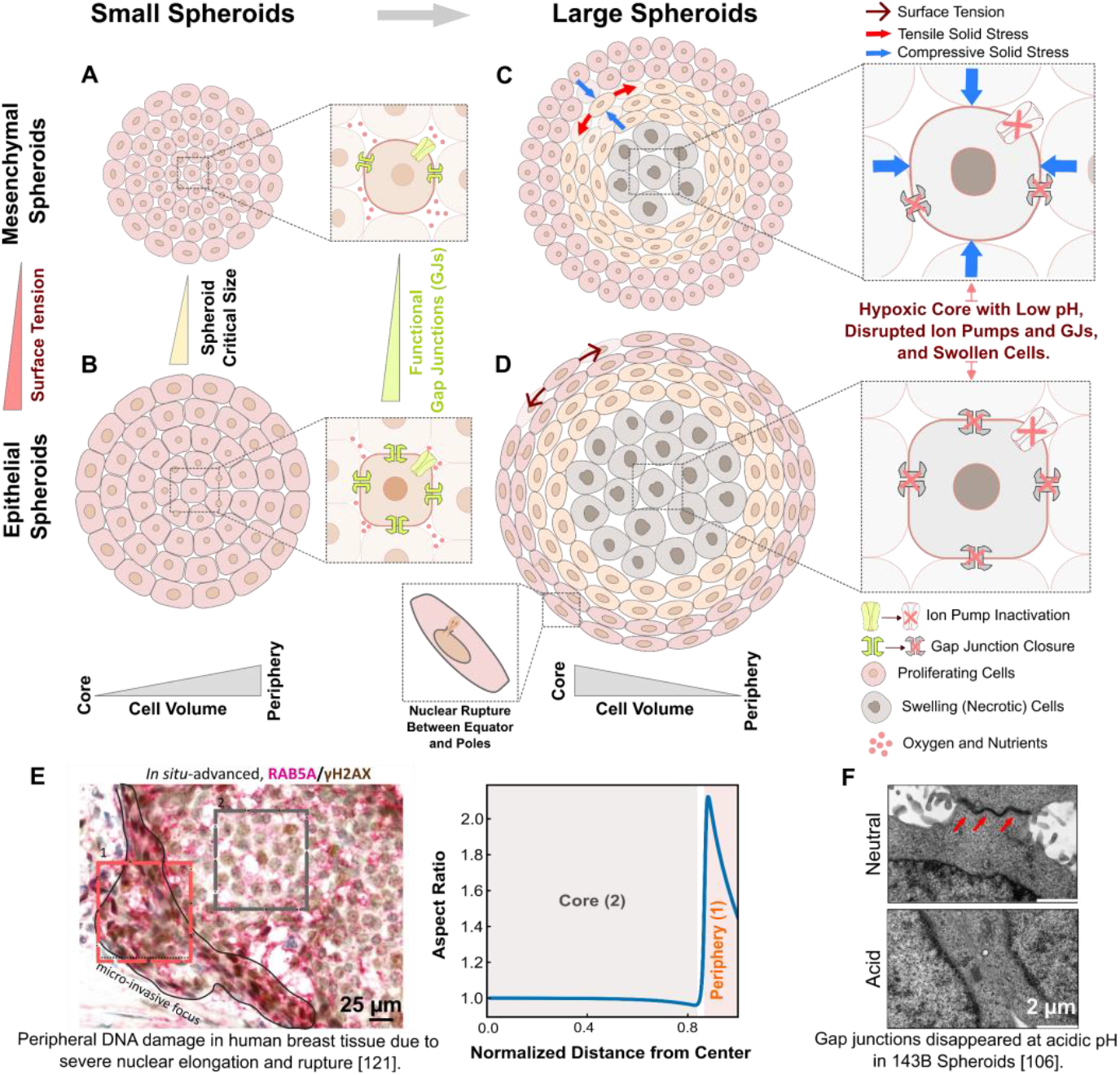
Metabolically driven core swelling regulates residual solid stress profiles and induces pronounced cell elongation in large tumor spheroids, which in turn promotes metastatic phenotypes through genomic instability and DNA damage. In schematic panels (**A–D**), nuclear shape is illustrated to approximately follow cell shape for clarity. **A)** In our previous work [31], we showed that in small spheroids with negligible metabolic gradients, fluid and ion transfer through gap junctions (GJs) increases cell volume from the core to the periphery. **B)** Experiments [109] have shown that epithelial tumor spheroids can use their extensive GJ networks to support transcellular nutrient transport. This reduces nutrient gradients and enables spheroids to grow larger without developing core hypoxia or necrosis. **C)** As spheroids continue to grow, metabolic gradients eventually emerge, leading to a hypoxic and acidic core. These conditions disrupt energy-intensive ion pumps and cause GJ closure, resulting in swelling of core cells. This central swelling generates isotropic compressive residual solid stress in the core and anisotropic residual stress at the periphery, with radial compression and tangential tension. Uniform osmotic swelling in the core also modulates cell morphology throughout the spheroid. Cells remain round at the core, become significantly elongated in the intermediate region, and gradually return to a more rounded shape toward the periphery. **D**) Cell shapes at the boundary of epithelial spheroids are nevertheless significantly elongated due to strong interfacial tensile forces in this region. **E)** This spatial pattern of cell morphology, with round cells at the core and elongated cells near the core and boundary, has also been observed in epithelial human breast tissues [121], consistent with model predictions. **F)** Transmission electron microscopy of 143B-derived tumor spheroids [106] shows disappearance of GJs at acidic pH.

While constrained cell proliferation has long been viewed as the primary driver of solid stress generation in tumor aggregates, the potential role of osmotic cell swelling has remained surprisingly unexplored. To our knowledge, this study presents the first theoretical framework, validated by experiments, that systematically investigates the contribution of osmotic swelling to solid stress generation in tumor spheroids. Using this framework, we first tested whether peripheral proliferation alone could explain the experimentally observed stress distributions. Analysis with proliferation as the sole source of volume expansion predicted peak compressive stress near the spheroid boundary (**Figure 3B**). However, this result contradicts multiple experimental studies [85-87], which consistently report maximum compressive stress localized at the spheroid core. Incorporating osmotic swelling into the model resolved this discrepancy by shifting the stress peak to the core region (**Figure 3F**), thereby aligning our predictions with experimental observations. These findings demonstrate that peripheral proliferation alone cannot explain the elevated compression observed at the spheroid core. Instead, core swelling emerges as the primary driver of this central compression. As the core swells, it exerts contact stresses on adjacent cell layers, generating strong radial compressive forces at their interface that propagate throughout the spheroid. Within the core, these forces produce large, uniform, and isotropic compression. Outside the core, however, the outward propagation of these compressive forces stretches successive cell layers tangentially, generating tangential tension. Taken together, our model predicts a stress distribution characterized by high compression in the spheroid core, balanced by tangential tension in the outer layers (**Figures 3F** and **6C**). Importantly, this predicted stress profile matches observations from cutting experiments performed on large tumor spheroids derived from HCT116 colon carcinoma cells [23, 24]. Partial cuts made into these spheroids consistently result in simultaneous bulging at the spheroid center and retraction at its boundary, revealing central compression and peripheral tangential tension. Notably, a similar compression-tension profile has been observed in brain, colon, breast and pancreatic tumor tissues ex vivo [15, 16], and in breast tumors in vivo [32], supporting the notion that in vitro spheroids faithfully recapitulate key biomechanical aspects of the tumor microenvironment.

Our model predicts a distinctive spatial pattern of cell shape within large tumor spheroids. Cells are nearly round in the necrotic core due to isotropic osmotic swelling, become markedly elongated in intermediate zones where stress anisotropy is high, and gradually return to more isotropic shapes toward the periphery as stress becomes less directional. At the boundary, cells remain minimally elongated in mesenchymal spheroids, where interfacial forces are weak (**Figures 6C** and **D**). By contrast, in epithelial spheroids, previous experiments have shown that cells become strongly elongated in the tangential direction, driven by pronounced polarization of cortical tension and large interfacial forces. To test these predictions, we compared them with published CT26 tumor spheroid observations [81] and performed independent validation in breast cancer systems, both in vitro using MDA-MB-231 spheroids and in patient-derived DCIS tissue sections. Using quantitative image analysis across spheroids and DCIS cross-sections spanning multiple growth stages, we quantified the spatial evolution of cell size and shape. The resulting morphology patterns closely follow theoretical predictions (**Figures 4 and 5**), supporting the robustness of our framework and its utility for interpreting clinically accessible tumor histology through a mechanobiological lens. The observed spatial organization of cell shapes, particularly the pronounced elongation in intermediate zones, further supports a central role for osmotic swelling in shaping the mechanical environment. Our model shows that reproducing this shape pattern requires incorporating core osmotic swelling, underscoring its essential role not only in generating stress profiles but also in driving spatial variations in cell morphology across the spheroid. A valuable insight from our theory is that cell shape serves as a practical, informative marker of local solid stress anisotropy within large spheroids. Our model quantitatively confirms this direct relationship: where anisotropy peaks, cell elongation is most pronounced, while isotropic stresses yield nearly spherical cell morphologies (**Figures 4D** and **5B**). Thus, cell shape functions as an effective, non-invasive, and spatially resolved indicator of the underlying stress environment, offering a valuable experimental and diagnostic readout. Furthermore, our findings extend beyond the initial metabolically driven swelling scenario. Even after extensive lysis of necrotic cells in the spheroid core, our theory predicts that the stress and morphological profiles remain largely unchanged (**Figure 5D**). This persistence stems from the Donnan osmotic effect, wherein negatively charged macromolecules released by lysed cells remain trapped within the core, sustaining its swollen, gel-like state (**Figure 5E**). As a result, elevated core osmotic pressure and, consequently, the associated stress distribution are maintained. Together, these insights establish a comprehensive and experimentally validated framework linking metabolic and ionic disruptions at the cellular scale to mechanical heterogeneity observed at the tissue level across tumor spheroids of varying growth stages. By leveraging cell morphology as a direct reporter of solid stress anisotropy, we provide an accessible method for inferring internal stress states, offering valuable implications for tumor diagnostics and mechanobiological research.

In our previous work [31], we demonstrated that fluid and ion transfer through gap junctions (GJs) increases cell volume from the core to the periphery in small spheroids (diameter∼75 *μm*) derived from MCF10A cells with negligible metabolic gradients (**Figure 6A**). However, our current theoretical and experimental findings reveal the opposite trend in large tumor spheroids with pronounced metabolic gradients: in these spheroids, cell volume increases from the periphery toward the core. This reversal arises from how GJ activity changes with spheroid growth. As spheroids enlarge, rapid oxygen depletion from the surface inward leads to the development of a hypoxic core. Cells in this region primarily rely on anaerobic glycolysis, resulting in lactic acid accumulation and a drop in both intra- and extracellular pH [42]. Typical pH values in the necrotic core of tumor spheroids (derived from brain, colon, and liver cancer cells) range from 6.6 to 7.0 [42-46], indicating a markedly acidic environment. Multiple experimental studies [36-40] have shown that under such acidic conditions, most gap junction channels close or disassemble, disrupting intercellular communication. Indeed, transmission electron microscopy of 143B-derived tumor spheroids directly visualized this dysfunction (**Figure 6F**), revealing complete loss of gap junctions under acidic conditions [106]. These findings show that necrotic core formation in tumor spheroids is accompanied by a loss of GJ-mediated communication, primarily due to acidification of the core microenvironment. Thus, while fluid flow through gap junctions increases cell volume toward the periphery in small spheroids, the breakdown of these channels in large spheroids, driven by metabolic dysregulation, along with the onset of osmotic swelling in the core, reverses this trend, resulting in increased cell volume in the core instead (**Figures 6C** and **D**). We verified this cell-volume reversal trend in our MDA-MB-231 tumor spheroids (**Figure S10**). Image analysis of the mid-plane of 4-day-old spheroids (**Figure S9**, radius ≈ 150 μm) revealed that these smaller spheroids, in contrast to larger 11-day-old spheroids, display an increase in cell area from the core toward the periphery. Therefore, the theory we presented in this work complements our previous model for small spheroids [31], providing a unified framework to understand spatial cell volume changes across all stages of spheroid growth. Future work could integrate these two models by explicitly considering the dependency of gap junction activity on local microenvironmental acidity. Interestingly, such an explicit relation has been experimentally obtained, where gap junction conductivity is a simple and sensitive function of intracellular pH, well-fitted by a Hill-type relation [39, 40].

The critical size at which a tumor spheroid develops severe core hypoxia, osmotic swelling, and necrosis, and is thus classified as *large*, depends primarily on the oxygen consumption rate (OCR) of its constituent cells, the diffusion coefficients of oxygen and other nutrients within the spheroid, and the duration of nutrient deprivation in the core. Cells with higher metabolic activity (i.e., higher OCR) consume oxygen more rapidly, generating steeper oxygen gradients (see **Eq. (14)**) and accelerating hypoxic core formation. Experiments analyzed in **Figures 4** and **5** show that spheroids derived from CT26 cells, which exhibit a higher OCR [107], display pronounced core swelling at smaller radii (∼120 *μm* versus ∼200 *μm*) compared to those derived from MDA-MB-231 cells, which have lower OCRs [108]. Oxygen availability in the spheroid core is also dictated by its internal microstructure. Oxygen and nutrients diffuse inward primarily through intercellular spaces, the extent of which depends on cellular phenotype. Epithelial spheroids feature densely packed cells (**Figures 6B** and **D**) with extensive (E-cadherin–mediated) junctions and minimal spacing, while mesenchymal spheroids feature looser architecture (**Figures 6A** and **C**) with increased ECM content. Notably, recent experiments [109] show that epithelial colon cancer spheroids (derived from HCT116 and HT29) employ gap junctions to enable transcellular nutrient transport and sustain core cells. This suggests that epithelial spheroids can harness gap junction networks as alternative diffusion routes, mitigating nutrient gradients and delaying hypoxia and necrosis (**Figure 6B**). Supporting this, fluorescence imaging of MCF-10A (epithelial) and MDA-MB-436 (mesenchymal) spheroids [82] reveals opposite cell volume trends despite almost equal size. In MCF-10A, volume increases from the center to the periphery, indicating functional gap junctions, while in MDA-MB-436, volume increases moving toward the core, suggesting central hypoxia. The duration of nutrient deprivation is another factor that determines if core conditions become severe enough to trigger extensive hypoxia and cell death. These hostile conditions occur when chronic nutrient starvation surpasses a critical threshold, depleting metabolic reservoirs and overwhelming adaptive mechanisms like gap junctions, ultimately disrupting ion homeostasis. Thus, prolonged deprivation is also essential for forming hypoxic and necrotic cores. This is supported by recent experimental findings [79] showing necrotic cores in day-20 T47D spheroids but not in size-matched day-5 spheroids. These spheroids were otherwise identical but generated using different initial seeding densities (4,000 cells/well for day 5 vs. 1,000 cells/well for day 20). Collectively, these findings highlight that the emergence of hypoxic and necrotic cores is a complex, dynamic process influenced by cell type, phenotype, and spheroid formation method. A thorough understanding of these factors is crucial for developing accurate 3D tumor models that closely replicate the in vivo tumor microenvironment, particularly regarding oxygenation status.

Experimental observations [110-112] reveal that tumor spheroids exhibit viscoelastic behavior arising from dynamic processes such as remodeling of structural proteins (e.g., actin filaments), adhesion (cadherin and integrin) turnover, and the movement and rearrangement of constituent cells. These dynamic processes, particularly cell motility, can potentially relieve solid stresses generated by metabolic dysregulation within the spheroid. However, experimental evidence suggests that these stresses do not fully relax, primarily due to continuous spheroid growth and subsequent structural changes. Cell movement and associated stress relief in spheroids are very slow, taking several hours for a cell to move its own diameter [82], comparable to the timescale of spheroid growth [113] and metabolic changes (approximately a day). As a result, although cellular rearrangements partially relax solid stresses, peripheral proliferation and core swelling continually regenerate stress, maintaining elevated stress levels. New growth-induced stress accumulates before previous stresses can fully dissipate. Importantly, cutting experiments [23] have directly shown that tumor spheroids accumulate solid stress over days of growth, confirming a build-up of solid stress over time. Additionally, as spheroids enlarge, internal cell movement and tissue fluidization become increasingly constrained, resulting in the accumulation of trapped solid stresses. In large spheroids, with pronounced metabolic gradients, interior cells become substantially immobilized due to local compaction and metabolic starvation, leading to a glassy or jammed state that significantly impedes stress relaxation within the spheroid core. The formation of this solid-like/jammed core is supported by theoretical models (e.g., vertex model [114]) as well as experimental observations [115]. Interestingly, even in small spheroids, where cells rearrange relatively freely, solid stress does not fully dissipate. This persistence is evidenced by differential cell swelling, caused by supracellular fluid flow through gap junctions driven by these stresses [31, 35]. Therefore, despite the viscoelastic nature of tumor spheroids, solid stresses do not fully dissipate within these cell aggregates and instead accumulate over time. Our solid elastic model, which neglects cell rearrangements and other dynamic processes, effectively captures this long-term stress profile. Future extensions could integrate these dynamics into our framework to provide deeper insights into stress evolution during key biological events such as epithelial-to-mesenchymal transition and tumor invasion.

Our findings are significant in the context of emerging recognition that solid stress has critical implications for cancer progression and metastasis. The mechanical environment within tumor spheroids and tissues, characterized by elevated compressive stresses and marked cell elongation, significantly impacts various biological processes that collectively enhance metastatic potential. Elevated compressive stresses, for example, have been experimentally linked to cell-cycle arrest in spheroids derived from human breast and colon cancer cells [116]. Our model’s predictions of high-stress environments thus offer important mechanistic insights into how these conditions may influence cellular proliferation. Additionally, the pronounced elongation observed in intermediate spheroid regions has profound consequences for mitotic fidelity. Flattened cells struggle to achieve proper mitotic rounding, disrupting spindle geometry and assembly [117]. Such disruptions can cause chromosome segregation errors, potentially leading to genomic instability and aneuploidy phenotypes commonly associated with aggressive cancer behavior [118]. Moreover, compressive stresses that diminish proliferation rates have also been shown to reduce the efficacy of chemotherapeutic agents targeting actively dividing cells, as demonstrated in pancreatic cancer spheroids [6]. Beyond mitotic fidelity, mechanical deformation due to cell elongation influences nuclear structure and mechanotransduction pathways critical for metastasis. Stretched nuclei exhibit expanded nuclear pores, facilitating the nuclear entry of transcriptional regulators such as YAP/TAZ (Yes-associated protein/transcriptional co-activator with PDZ-binding motif) [119]. Once inside, YAP/TAZ promotes transcriptional programs that enhance migration, invasion, and metastatic capabilities, and has been linked to resistance mechanisms across chemotherapy, targeted therapy, and immunotherapy [120]. Immunohistochemical analysis of advanced in situ human breast tumors (**Figure 6E**) further shows that extreme cellular elongation at the tumor periphery can compromise nuclear integrity, leading to rupture of the nuclear envelope [121]. Our model provides a mechanistic explanation for this rupture by predicting how nuclear envelope tension and Gaussian curvature vary across elongated, oblate nuclei (**Supplementary Note S7**). Specifically, while tension is greatest near the poles and curvature peaks at the equator, their opposing spatial distributions create a structurally vulnerable zone between these regions where nuclear rupture is most likely to occur (**Figures 6D** and **S4**), consistent with experimental observations [121]. Such rupture events allow cytoplasmic factors, including the ER-associated exonuclease TREX1, to enter the nucleus and induce DNA damage, thereby fostering invasive phenotypes [116]. Expanding our mechano-electro-osmotic model to incorporate these complex mechanobiological interactions could provide deeper insights into tumor spheroid biology, serving as robust in vitro mimics of native tumor conditions. Taken together, these results indicate that spatially heterogeneous solid stresses and deformation states in the tumor microenvironment can drive multiple hallmarks of aggressive behavior, including altered proliferation, therapy resistance, YAP/TAZ activation, and nuclear damage. A major next step is to link these mechanical cues to the molecular programs they induce, motivating the integration of our model predictions with spatially resolved gene-expression measurements.

Building on this motivation, the mechanical insights provided by our model, including spatial variations in cell morphology and the solid stress field, can be integrated with spatial transcriptomics to uncover the molecular programs underlying cancer progression and invasion in multicellular aggregates. Such integrative approaches have recently proven powerful in melanoma [122], where spatial transcriptomics revealed that mechanical confinement imposed by the surrounding microenvironment can epigenetically reprogram tumor cells at the tumor– tissue interface, inducing stable chromatin remodeling and a transition from a proliferative to an invasive, neuron-like transcriptional state. Motivated by these findings, a natural extension of our work is the joint analysis of transcriptional and mechanical signals in tumor spheroids. Importantly, recent methodological advances [123, 124] now enable rapid, high-resolution spatial transcriptomic profiling in spheroid systems. Using previously published spatially resolved single-cell RNA sequencing data, we performed a preliminary analysis of large MDA-MB-231 tumor spheroids [119] (**Supplementary Note S8**). This analysis reveals a clear correlation between the core–periphery solid-stress landscape predicted by our model and spatial patterns of gene expression. In particular, the expression of tension-sensitive genes (**Figure S5B**) and YAP target genes (Figure **S5C** and **S5D**) increases from the spheroid core toward the periphery, consistent with the predicted gradients in solid stress and cell morphology. A more comprehensive dissection of the underlying mechanotransduction pathways will require future studies combining additional multi-omics measurements with carefully designed mechano-chemical perturbation experiments.

In summary, by highlighting osmotic cell swelling, driven by metabolic dysregulation, as a fundamental mechanism underlying residual solid stress generation, our findings establish crucial connections between metabolic dysfunction, cellular deformation, and metastatic progression. These insights pave the way for novel mechanotherapeutic strategies aimed at alleviating pathological stress within the tumor microenvironment.

## Methods

### 1. Spheroid Model Generation

Invasive breast cancer MDA-MB-231 cells (gift from Philippe Chavrier’s laboratory) were cultured in high-glucose DMEM medium (Catalog No. MT10-013-CV) supplemented with 10% fetal bovine serum and 1% penicillin–streptomycin (Catalog No. 15140122). For spheroid generation, cells were detached using TrypLE™ Express (Catalog No. 12605-010) and seeded at a density of 1,000–2,000 cells per well in Nunclon™ Sphera 96-well round-bottom ultra-low attachment plates (Catalog No. 174929) to form compact spheroids. Spheroids were cultured for 4, 11, and 14 days to model progressive stages of necrosis and then embedded in 4 mg mL^-1^ Type I collagen (Catalog No. 354249).

### 2. Immunofluorescence Staining & Confocal Microscopy

Collagen gels with embedded cancer cell spheroids were fixed with 4% paraformaldehyde at room temperature for 2 h. To increase cell membrane permeability for antibody staining, the gels were incubated in PBS containing 2% Triton X-100 for 48 h. The gels were then blocked in PBS containing 10% goat serum, 0.5% Triton X-100, and 2.5% DMSO (blocking buffer) for 6 h and subsequently incubated with primary antibody against Ki67 (1:500; Abcam #ab15580) diluted in blocking buffer for 24 h. The gels were then incubated with the corresponding secondary antibody, along with phalloidin (1:200; Thermo Fisher Scientific #A12380) and SYTOX (1:1500; Thermo Fisher Scientific #S11380), for 24 h. Finally, the gels were immersed in RapiClear™ 1.49 clearing solution (SunJin Lab; RC149001) for 24 h to enhance light penetration for Airyscan confocal microscopy (Zeiss LSM 900). All immunofluorescence staining steps were performed at 4 °C with slow continuous shaking.

### 3. Tumor Tissue Acquisition and Staining

De-identified human breast tumor specimens were obtained with institutional review board (IRB) approval from the University of Pennsylvania. It was determined that the proposal meets eligibility criteria for IRB review exemption authorized by 45 CFR 46.104, category (IRB PROTOCOL#: 856666). Briefly, tumor tissues were fixed with 10% neutral buffered formalin, dehydrated in ethanol, and paraffin embedded using standard protocols. Subsequently, FFPE tissue blocks were deparaffinized and 5 μm sections were stained with Hematoxylin & Eosin (Dako, CS70030-2) to visualize nuclei and overall tissue morphology. The slides were then manually screened and annotated for necrosis by a pathologist and imaged using brightfield microscopy (Zeiss; Axio Vert.A1).

### 4. Image Analysis

#### AI-based Cell Segmentation using the CellSAM Model

Cell segmentation was performed using an artificial intelligence–based pipeline built on the CellSAM model [92], a deep-learning framework designed for high-precision cellular segmentation in fluorescence microscopy. In our analysis, both the nuclear (Sytox) channel and the cell-boundary (phalloidin) channel were jointly used as inputs to CellSAM, enabling the model to integrate complementary structural information when identifying individual cells. By leveraging nuclear localization for cell separation and phalloidin-labeled membranes for boundary definition, this dual-channel strategy ensures robust segmentation that avoids both over-segmentation (splitting one cell into multiple fragments) and under-segmentation (merging adjacent cells into a single object). CellSAM, which adapts the Segment Anything Model (SAM) architecture for biological imaging, was tuned for our dataset of 2D middle-plane confocal slices of spheroids, providing accurate and biologically consistent cell masks across heterogeneous tissue regions.

#### Image Preprocessing and CellSAM Segmentation

Raw experimental images were first imported and preprocessed using custom Python scripts leveraging the SimpleITK library. Images were converted to grayscale, normalized to enhance contrast, and formatted as 8-bit RGB arrays compatible with the CellSAM model input requirements. The segmentation pipeline produced binary masks delineating each identified cell within the images. These masks served as the basis for subsequent morphological analyses.

#### Post-processing and Ellipse Fitting Analysis

Segmented masks were imported into MATLAB using custom-developed scripts. To refine segmentation outcomes, interactive mask editing was performed in MATLAB, allowing the removal of spurious or overlapping masks through manual validation. Following mask refinement, MATLAB’s *regionprops* function was utilized to fit ellipses to the masks, providing precise measurements of cell dimensions and orientations. From these ellipse fits, centroids, sizes, and aspect ratios of individual cells were determined.

#### Spatial Binning and Trend Analysis

To capture spatial morphological gradients, segmented cells were binned radially from the spheroid center to the periphery, aligning bins with cellular layers observed within the spheroid structure. Cell properties such as cross-sectional area and aspect ratio were averaged within each radial bin, and statistical error was computed as the standard error of the mean. This binning approach facilitated clear visualization and quantification of morphological trends across the spheroid radius.

#### Validation and Visualization

Quantitative outcomes of segmentation and ellipse fitting were validated visually by overlaying ellipse contours onto original microscopy images using MATLAB visualization tools. Additionally, comparative analyses between computational predictions and experimental observations were visualized through custom plots, illustrating trends in cell morphology against radial position.

### 5. Reanalysis of Diffusion Smart-seq3D Spheroid Transcriptomes

We reanalyzed the published diffusion Smart-seq3D single-cell transcriptomics dataset from MDA-MB-231 tumor spheroids [124]. In the original study, each cell is assigned a continuous radial coordinate *r* by log-transforming Calcein-AM fluorescence and rescaling to [0, 1], where *r* = 0 denotes the spheroid core and *r* = 1 the periphery; we used the same radial coordinate definition in our analysis. For each cell and each gene set, we computed the gene-set UMI count *y*_set,*i*_ as the sum of UMIs across genes in the set, and the total UMI count *N*_total,*i*_ as the total UMIs across all genes in that cell. Gene-set activity was quantified as gene-set fractional abundance *f*_set,*i*_ = *y*_set,*i*_/*N*_total,*i*_. We analyzed four programs: MSigDB HALLMARK_HYPOXIA, MSigDB CORDENONSI_YAP_CONSERVED_SIGNATURE, a tensile-responsive gene set derived *a priori* from an independent cyclic tensile strain RNA-seq dataset by selecting genes that increased strictly monotonically with strain magnitude [125] (**Table S2**), and a curated canonical YAP target mini-set (**Table S3**).

To test for spatial regulation along the continuous radial axis, we fit negative binomial generalized linear models to the gene-set counts *y*_set,*i*_ with an offset for sequencing depth log (*N*_total,*i*_). Following the original Smart-seq3D analysis framework [124], we used a quadratic dependence on radial position to capture potentially non-monotone core–periphery trends. Specifically, we modeled the expected mean gene-set count *μ*_*i*_ = 𝔼[*y*_set,*i*_] using a log link, 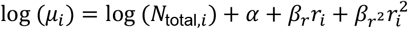, and compared this quadratic model to a null model that included only the offset and intercept (log (*μ*_*i*_) = log (*N*_total,*i*_) + α). Model significance was assessed by a likelihood ratio test. Model effect size was summarized as the periphery-to-core fold change, defined as the ratio of the model-predicted mean at *r* = 1 to that at *r* = 0, that is *μ*(1)/*μ*(0); under the quadratic model above, 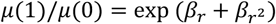. As an additional nonparametric check, we computed the Xi correlation p-value between gene-set fractional abundance *f*_set,*i*_ and radial position *r*_*i*_. For hypoxia, we also performed a pre-specified core-versus-rim comparison using the bottom 20% versus top 20% of the *r* distribution and assessed differences in *f*_set,*i*_ using a two-sided Wilcoxon rank-sum test.

For visualization, we plotted single-cell fractional abundance versus *r* (grey points), overlaid binned means computed in 10 radial bins (red circles) with error bars reflecting uncertainty of the binned mean, and superimposed the fitted negative binomial quadratic trend as a black curve.

## Data availability

Raw images from in vitro spheroid experiments and patient-derived tissue sections are available from the corresponding author upon reasonable request. All experimental measurements and simulation outputs supporting the figures and conclusions of this study are provided in the accompanying Source Data file.

## Code availability

Mathematica and R code to reproduce **Figures 5B** and **S5B** are available on GitHub: https://github.com/MohammadDehghany/Tumor-Spheroid-Model.

Code used for segmentation and quantification of experimental images is publicly available at: https://github.com/vs-vivek/Tumor_Spheroid_Analysis.

## Acknowledgements

This work was supported by NCI Award U54CA261694 (V.B.S.); NSF CEMB Grant CMMI-154857 (V.B.S.); NSF Grant DMS-2347834 (V.B.S.); NIBIB Awards R01EB017753 (V.B.S) and R01EB030876 (V.B.S.) and NIGMS Award R01GM155943 (V.B.S).

