## Supplementary Material for "Metabolic starvation–induced cell swelling drives solid stress in tumors"

Supplementary information for:

### Metabolic starvation–induced cell swelling drives solid stress in tumors

**Supplementary Note S1:** *Cortical stress and strain in deformed cells.*

Consider a spherical cell with radius $a_0$ that deforms into an oblate spheroid with semi-axis lengths $a(r)$ along the $z$-axis and $b(r)$ along the $x$ and $y$-axes at radial position $r$ within the tumor spheroid (**Figure S1**, upper panel). Owing to the symmetry of deformation, the cell cortex geometry can be fully described by a single meridional profile (**Figure S1**, lower panel). Let point $\mathrm{P}$, with coordinates $(X, Z)$ or $(S, \Psi)$ on the relaxed cortex, move to point $\mathrm{Q}$, with coordinates $(x, z)$ or $(s, \psi)$ on the deformed cell. Here, $S$ and $s$ are the arc lengths measured along the meridian from the pole to $\mathrm{P}$ and $\mathrm{Q}$, respectively, and $\Psi$ and $\psi$ are the corresponding polar angles.

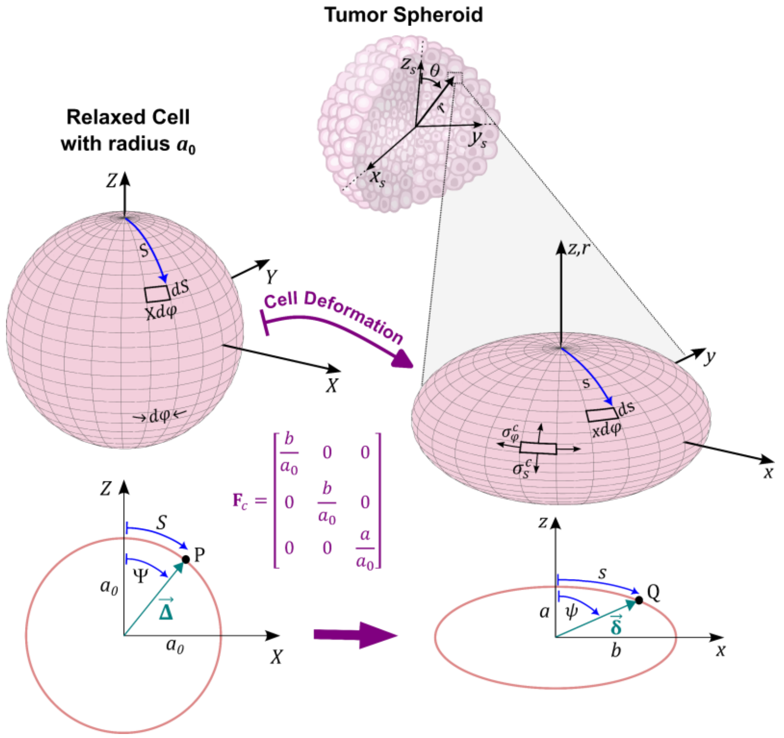


**Figure S1: Geometric description of cell deformation from a relaxed sphere to an oblate spheroid within a tumor spheroid.** Schematic illustrating the transformation of a relaxed spherical cell (radius $a_0$) into an oblate spheroid with semi-axis lengths $a(r)$ (along the $z$-axis) and $b(r)$ (along the $x$ and $y$-axes) at radial position $r$ in the tumor spheroid. The relaxed and deformed cell shapes are described by their meridional profiles, parameterized by arc lengths ($S$ and $s$) and polar angles ($\Psi$ and $\psi$). The deformation gradient tensor $\mathbf{F}_c$ relates the relaxed ($\vec{\mathbf{\Delta}}$) and deformed ($\vec{\boldsymbol{\delta}}$) position vectors. The lower panels illustrate the projection of the 3D geometry onto the meridional plane. This framework is used to compute cortical strains and stresses in the hoop ($\sigma_\varphi^c(r,s)$) and meridional ($\sigma_s^c(r,s)$), directions.

This deformation induces cortical stresses in the cell membrane, denoted by $\sigma_\varphi^c(r,s)$ and $\sigma_s^c(r,s)$, acting along the hoop ($\varphi$) and meridional ($s$) directions, respectively. To determine these stress components, we first calculate the associated stretch fields:

$$\lambda_\varphi(r,s) = \frac{xd\varphi}{Xd\varphi} = \frac{x}{X} \tag{S1}$$

$$\lambda_s(r,s) = \frac{ds}{dS} \tag{S2}$$

where $\lambda_\varphi$ and $\lambda_s$ are the hoop and meridional stretches. To evaluate these stretches, we represent point $\mathrm{P}$ in the relaxed configuration by position vector $\vec{\boldsymbol{\Delta}} = a_0(\sin(\Psi), 0, \cos(\Psi))$ and point $\mathrm{Q}$ in the deformed configuration by $\vec{\boldsymbol{\delta}} = \delta(r)(\sin(\psi), 0, \cos(\psi))$. These vectors are related by the cell deformation gradient tensor $\mathbf{F}_c = a_0^{-1}\begin{bmatrix} b(r) & 0 & 0 \\ 0 & b(r) & 0 \\ 0 & 0 & a(r) \end{bmatrix}$ through the relation:

$$\vec{\boldsymbol{\delta}} = \mathbf{F}_c\vec{\boldsymbol{\Delta}} \tag{S3}$$

From this relation, we obtain:

$$x = \delta(r)\sin(\psi) = b(r)\sin(\Psi), \tag{S4}$$

$$z = \delta(r)\cos(\psi) = a(r)\cos(\Psi). \tag{S5}$$

Using these relations, and noting that $X = a_0\sin(\Psi)$, $dS = a_0 d\Psi$, and $ds = \sqrt{dx^2 + dz^2}$ (see **Figure S1**), the stretches in the hoop and meridional directions can be obtained as:

$$\lambda_\varphi(r) = \frac{x}{X} = \frac{b(r)}{a_0} \tag{S6}$$

$$\lambda_s(r,\psi) = \frac{ds}{dS} = \frac{a(r)}{a_0}\sqrt{\frac{b^4(r) + \tan^2(\psi)\,a^4(r)}{a^2(r)b^2(r) + \tan^2(\psi)\,a^4(r)}} \tag{S7}$$

These results indicate that the hoop stretch remains uniform across the membrane of each cell, whereas the meridional stretch varies along the membrane, decreasing from the pole ($\psi = 0$) to the equator ($\psi = \pi/2$). From these stretches, the corresponding Lagrangian (Green) strain fields can be then calculated as:

$$\varepsilon_\varphi(r) = 0.5\left(\lambda_\varphi^2(r) - 1\right) \tag{S8}$$

$$\varepsilon_s(r,\psi) = 0.5\left(\lambda_s^2(r,\psi) - 1\right) \tag{S9}$$

To obtain the stress-strain relationship for the cell cortex, we next examine its stress state in more detail. Since the cortex thickness is much smaller than the cell dimensions ($h/a, h/b \ll 1$) [1], stresses perpendicular to the cortex, namely solid stresses and hydrostatic pressures, are significantly smaller than the in-plane cortical stresses (see **Eqs. (S12)** and **(S13)** and **Figure S2**). As a result, the cortex can be approximated as being in a plane stress state. Under these conditions, it can be proven [2] that, for any arbitrary large deformation, the stress-strain law for the cell cortex can be expressed as [2]:

$$\sigma_\varphi^c(r,\psi) = \frac{E}{1-\nu^2}\left(\varepsilon_\varphi^c(r) + \nu\varepsilon_s^c(r,\psi)\right) + \sigma_\varphi^{act}(r), \tag{S10}$$

$$\sigma_s^c(r,\psi) = \frac{E}{1-\nu^2}\left(\varepsilon_s^c(\mathrm{r},\psi) + \nu\varepsilon_\varphi^c(r)\right) + \sigma_s^{act}(r). \tag{S11}$$

Here, the second terms $\sigma_{s(\varphi)}^{act}$ represent the active stresses generated by myosin motors in the cortex, while the first terms correspond to elastic cortical stresses, which are formally analogous to the familiar linear elasticity law for plane-stress conditions. $E$ and $\nu$ denote the elastic material properties of the cortex, which generally depend on the strain invariants [2]. In our analysis, we assume the cortex to be linearly elastic, so that $E$ and $\nu$ are taken as constants.

To relate the cortical stresses derived above to the solid stress components, $\sigma_r$ and $\sigma_\theta$, as well as to the hydrostatic pressure difference $\Delta P$ across the cell membrane, we next consider the mechanical equilibrium of the cell in the spheroid's radial and tangential directions (**Figure S2**). The corresponding balance equations are:

$$\sigma_r(r) = \frac{2h\sigma_s^c\left(r, \psi = \frac{\pi}{2}\right)}{b(r)} - \Delta P(r) \tag{S12}$$

$$\sigma_\theta(r) = \frac{4h\int_0^{\frac{\pi}{2}} \sigma_\varphi^c(r,\psi)\, ds}{\pi b(r) a(r)} - \Delta P(r) \tag{S13}$$

The cortical stresses in these expressions, obtained from **Eqs. (S6)–(S11)**, are:

$$\sigma_s^c\left(r, \psi = \frac{\pi}{2}\right) = \frac{(\nu b^2(r) + a^2(r) - (\nu+1)a_0^2)}{2(1-\nu^2)a_0^2} E + \sigma_s^{act}(\mathrm{r}) \tag{S14}$$

$$\int_0^{\frac{\pi}{2}} \sigma_\varphi^c(r,\psi) ds \approx \frac{\left(a^2(r) + 3b^2(r)\right)\pi}{8b(r)} \sigma_\varphi^{act}(\mathrm{r}) + \frac{\pi\left((2\nu+3)b^4(r) + (a^2(r) - 3a_0^2)(\nu+1)b^2(r) + a^2(r)\left((a^2(r) - a_0^2)\nu - a_0^2\right)\right)}{16a_0^2(1-\nu^2)b(r)} E \tag{S15}$$

In deriving **Eq. (S15)** from **Eq. (S13)**, truncated Legendre series were used to approximate the elliptic integrals that arise. We verified that these approximations introduce at most a 10% error for the aspect ratio range relevant to this study ($1 \le b(r)/a(r) \le 2$).

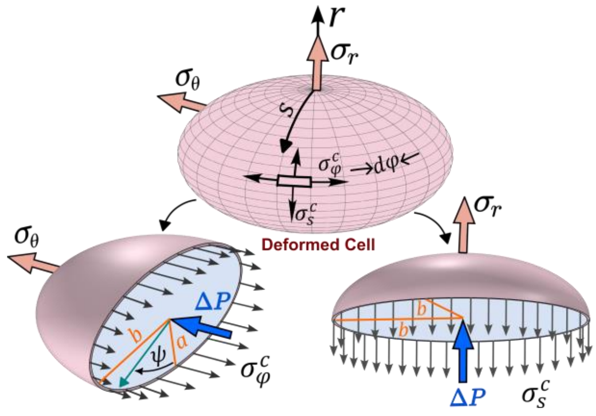


**Figure S2: Mechanical equilibrium of a deformed cell within a tumor spheroid.** Schematic of a deformed cell subjected to solid stresses ($\sigma_r, \sigma_\theta$), cortical stresses in the meridional ($\sigma_s^c$) and hoop ($\sigma_\varphi^c$) directions, and a hydrostatic pressure difference ($\Delta P$) across the membrane. The top panel illustrates the cortical stress directions on the cell surface, while the lower panels depict cross-sections highlighting the semi-axes, polar angle $\psi$, and the distribution of cortical stresses that maintain force balance in the radial (right) and tangential (left) directions

**Supplementary Note S2:** *Elastic and proliferative decomposition of spheroid growth and derivation of proliferative stretch*

As explained in the main text (Section 2.4), growth in large spheroids has two principal contributions: an elastic component, arising from changes in cell dimensions such as overall size and aspect ratio, and a proliferative component, associated with displacements due to cell division (see **Figure S3**). We describe these combined deformations using a multiplicative decomposition of the deformation gradient: $\boldsymbol{F} = \boldsymbol{F_e}\boldsymbol{F_p}$, where the total deformation gradient $\boldsymbol{F}$ is expressed as the product of elastic ($\boldsymbol{F_e}$) and proliferative ($\boldsymbol{F_p}$) parts. Taking the early-stage spheroid as the reference configuration (**Fig. 1D**), with the radial position of a cell denoted by $R$, we define the radial displacement during growth as $u_r(R) = r - R$. The radial and tangential components of the macroscopic stretch then follow as:

$$\begin{bmatrix} \frac{du_r(R)}{dR}+1 & 0 & 0 \\ 0 & \frac{u_r(R)}{R}+1 & 0 \\ 0 & 0 & \frac{u_r(R)}{R}+1 \end{bmatrix} = \begin{bmatrix} \frac{a(r)}{a_0} & 00 & 0 \\ 0 & \frac{b(r)}{a_0} & 0 \\ 0 & 0 & \frac{b(r)}{a_0} \end{bmatrix} \begin{bmatrix} \lambda_{p,r} & 0 & 0 \\ 0 & \lambda_{p,\theta} & 0 \\ 0 & 0 & \lambda_{p,\theta} \end{bmatrix} \tag{S16}$$

Here, we assume that the ratios $a(r)/a_0$ and $b(r)/a_0$ represent elastic stretches due to changes in cell morphology in the radial and tangential directions, respectively, while $\lambda_{p,r}$ and $\lambda_{p,\theta}$ capture proliferation-induced stretches in the corresponding directions. To estimate these stretches, we consider a single cell layer at radius $R$ in the reference configuration containing $n$ cells. During the first proliferation cycle, a fraction $\rho_{\theta,1}$ of these cells divide with cleavage planes oriented tangentially, thereby adding $\rho_{\theta,1}n$ new cells. The layer thus contains $n(1+\rho_{\theta,1})$ cells after this step and has radius $r_1$ (**Fig. S3**). The circumferential stretch associated with this growth step is then:

$$\frac{u_r(R)}{R}+1 = \frac{r_1}{R} = \frac{(1+\rho_{\theta,1})b(r_1)}{a_0} \tag{S17}$$

In the same manner, the tangential stretch in subsequent growth steps can be obtained. For instance, after the second proliferation cycle, $r_2/r_1 = (1+\rho_{\theta,2})b(r_2)/b(r_1)$, where $2b(r_2)$ is the cell length in the tangential direction after the second cycle and $\rho_{\theta,2}$ is the proliferating fraction. By multiplying contributions from successive cycles, the total tangential stretch accumulated over $m$ proliferation cycles is:

$$\frac{u_r(R)}{R}+1 = \frac{r}{R} = \frac{r_1}{R}\times\frac{r_2}{r_1}\times \dots \times\frac{r}{r_{m-1}} = \prod_{1}^{m}(1+\rho_{\theta,m})\frac{b(r)}{a_0} \tag{S18}$$

Here, $m$ denotes the number of proliferation cycles between the reference and current configurations. Comparing this result with **Eq. (S17)**, we identify the proliferative component of the tangential stretch as: $\lambda_{p,\theta} = \prod_1^m (1 + \rho_{\theta,m})$.

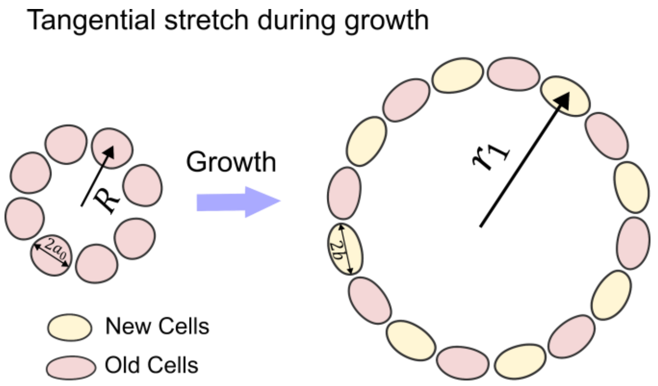


**Figure S3: Tangential stretch during the first proliferation cycle.** A single cell layer at radius $R$ in the reference configuration contains $n = 8$ cells. When a fraction $\rho_{\theta,1} = 1$ of these cells divide tangentially, $\rho_{\theta,1} n = 8$ new cells (shown in yellow) are added. This increases the layer radius to $r_1$, and the circumferential stretch is given by Eq. (S17).

For isotropic proliferation ($\lambda_{p,r} = \lambda_{p,\theta}$), the fraction of proliferating cells in each single direction can be at most one third. Under these conditions, higher-order terms can be neglected, and the proliferative stretch (to the first order) may be approximated as $\lambda_{p,\theta} \approx 1 + \sum_1^m \rho_{\theta,m}$. Experimental observations [3-5] indicate that $\rho_{\theta,m}$ is approximately uniform across the spheroid during early cycles and later decreases toward the core roughly exponentially. To capture this, we model the (isotropic) proliferative stretch as:

$$\lambda_{p,r} = \lambda_{p,\theta} = 1 + \rho_e + \rho_l e^{-(r_{out} - r)/(w)} \tag{S19}$$

where $\rho_e$ denotes the effective (accumulated) proliferating fraction during early cycles, and $\rho_l$ represents the proliferating fraction at the spheroid boundary during later cycles. The parameter $w$ is a characteristic decay length that sets how rapidly proliferation decreases toward the core.

**Supplementary Note S3:** *Cell stiffness anisotropy leads to physically inconsistent results*

Our results indicate that when spheroid cells are modeled as isotropically stiff, the compressive solid stresses generated by peripheral proliferation and interfacial forces decrease with depth and stabilize in the core (**Figure 3B**). This prediction contrasts with experimental findings [6-9], which show that compressive stresses increase toward the core. To reconcile this discrepancy, previous studies [9, 10] have proposed that cells within tumor spheroids exhibit stiffness anisotropy, with higher stiffness in the radial direction, thereby promoting stress accumulation at the core. However, we show that this assumption leads to inconsistent predictions and does not fully account for the experimentally observed stress profiles.

The stress–strain relationship for an anisotropic elastic sphere with spherical symmetry is given by [11]:

$$\varepsilon_\theta = \varepsilon_\phi = \frac{(1-\nu_1)}{E_\theta}\sigma_\theta - \frac{\nu_2}{E_r}\sigma_r, \tag{S20}$$

$$\varepsilon_r = \frac{1}{E_r}\sigma_r - \frac{2\nu_2}{E_\theta}\sigma_\theta, \tag{S21}$$

where $\nu_1$ and $\nu_2$ are Poisson's ratios, and $E_r$ and $E_\theta$ are the elastic moduli in the radial and tangential directions, respectively. The strain components can be written in terms of the radial displacement $u_r$ as $\varepsilon_\theta = u_r/r$ and $\varepsilon_r = du_r/dr$. Substituting these expressions into **Eqs. (S20)** and **(S21)**, and solving for the solid stress components, we obtain:

$$\sigma_r = \frac{\eta E_\theta\left((1-\nu_1)\frac{du_r}{dr} + 2\nu_1\frac{u_r}{r}\right)}{1-\nu_1-2\nu_1^2}, \tag{S22}$$

$$\sigma_\theta = \frac{E_\theta\left(\nu_1\frac{du_r}{dr} + \frac{u_r}{r}\right)}{1-\nu_1-2\nu_1^2}, \tag{S23}$$

where we have set $\nu_2 = \nu_1$ for simplicity and defined $E_r = \eta E_\theta$, with $\eta > 1$, to represent higher stiffness in the radial direction relative to the tangential direction. By substituting **Eqs. (S22)** and **(S23)** into the equilibrium equation in spherical coordinates (**Eq. (1)**) and solving for $u_r$, it can be shown that, after imposing the boundary condition $u_r(0) = 0$, the solid stress components take the form $\sigma_r \propto r^{-\beta}$ and $\sigma_\theta \propto r^{-\beta}$, where

$$\beta = \frac{\eta(3-\nu) - 2\nu - \sqrt{(9\nu^2 - 6\nu + 1)\eta^2 - 4\eta(\nu^2 + 3\nu - 2) + 4\nu^2}}{2\eta(1-\nu)}. \tag{S24}$$

It can be shown (see proof below) that $\beta$ is always positive for $0 < \nu \leq 0.5$ and $\eta > 1$, implying that both solid stress components diverge toward the center. As a result, these stresses become unbounded at the spheroid center, which is physically unrealistic. More importantly, from the equilibrium equation (**Eq. (1)**), we find that solid stress anisotropy, given by $\sigma_\theta - \sigma_r = \frac{r}{2}\frac{d\sigma_r}{dr} \propto r^{-\beta}$, also increases toward the center. This prediction contradicts recent experimental findings [6, 7], which reveal that stress within the spheroid core is nearly isotropic.

***Proof***: Since the denominator $2\eta(1-\nu)$ is always positive for $0 < \nu \leq 0.5$ and $\eta > 1$, it suffices to show that the numerator is also always positive under these conditions. To this end, we rewrite the numerator as: $P - \sqrt{Q}$, where $P = \eta(3-\nu) - 2\nu$, and $Q = (9\nu^2 - 6\nu + 1)\eta^2 - 4\eta(\nu^2 + 3\nu - 2) + 4\nu^2$. Since both $P > 0$ and $\sqrt{Q} > 0$, proving $P > \sqrt{Q}$ is equivalent to showing that $P^2 - Q > 0$. After some simplifications, we obtain: $P^2 - Q = 8\eta(\eta-1)(1-\nu^2) > 0$, which is strictly positive for all $0 < \nu \leq 0.5$ and $\eta > 1$. Therefore, both the numerator and the denominator of **Eq. (S24)** are always positive, and we conclude that $\beta > 0$ under the given conditions.

**Supplementary Note S4:** *Local stress anisotropy correlates with cell aspect ratio***.**

We next derive the relationship between the cell aspect ratio ($Ar = b/a$) and the solid stress anisotropy, defined as the difference between the circumferential and radial stresses ($\sigma_\theta - \sigma_r$) within the spheroid. Using **Eqs. (S12)–(S15)** and carrying out algebraic simplifications, the stress anisotropy as a function of the cell aspect ratio can be expressed as:

$$\sigma_\theta - \sigma_r = \frac{Eh}{2Vr(\nu^2 - 1)Ar^2 a_0}\left(\frac{Vr}{Ar^2}\right)^{\frac{1}{3}}\left((3Ar-1)(\nu+1)(\gamma(\nu-1)+0.5)Ar^2\left(\frac{Vr}{Ar^2}\right)^{\frac{1}{3}} - Vr\frac{(2\nu+3)Ar^3 + (3-2\nu)Ar^2 + (4-\nu)Ar - \nu}{2}\right)(Ar - 1) \quad \text{(S25)}$$

Here, we have assumed isotropic contractility, meaning $\sigma^{act} = \sigma_s^{act} = \sigma_\varphi^{act}$, and defined $\gamma = \sigma^{act}/E$. Additionally, $Vr = b^2 a/a_0^3$ denotes the volume ratio of the cell relative to its physiological volume. Equation **(S25)** shows that a spherical cell shape ($Ar = 1$) corresponds to an isotropic stress field ($\sigma_\theta = \sigma_r$) in the spheroid. To examine how cell shape and the solid stress field are related when the stress field is not isotropic, we next calculate the derivative of the stress anisotropy with respect to the aspect ratio:

$$\frac{d(\sigma_\theta - \sigma_r)}{dAr} = \frac{3Eh}{a_0 Ar^{11/3} Vr^{5/3}(1-\nu^2)}\left(\frac{(1-\nu^2)\left(\frac{Vr}{Ar^2}\right)^{\frac{4}{3}} Ar^4(3Ar^2+2Ar-2)\gamma}{9} + \frac{Vr^2\big((4\nu+6)Ar^4 - 2Ar^3\nu - (\nu+1)Ar^2 + 10Ar - 4\nu\big) - (\nu+1)\left(\frac{Vr}{Ar^2}\right)^{\frac{4}{3}} Ar^4(3Ar^2+2Ar-2)}{18}\right) \quad \text{(S26)}$$

Given that $0 < \nu \leq 0.5$, and noting that the cell aspect ratio and volume ratio are typically close to or greater than unity ($Ar \geq 1$, $Vr \geq 1$) in large tumor spheroids (**Figs. 4** and **5**), the first term of the derivative in **Eq. (S26)** is always positive (indeed, it remains positive for all $Ar \geq 0.55$). Furthermore, although the proof is straightforward yet lengthy and therefore omitted here, the second term can also be shown to be strictly positive and to increase with $Ar$ under these conditions. More generally, this positivity holds for all $Ar \geq 0.55$ and $Vr \geq 0.28$. Taken together, these results demonstrate that cells subjected to larger stress anisotropy exhibit more pronounced tangential elongation, thereby establishing a direct relationship between cell shape and stress anisotropy: $(\sigma_\theta - \sigma_r) \propto (Ar - 1)$.

**Supplementary Note S5:** *Material parameters and motivation.*

The spheroid radius $r_{out}$ in each case was directly measured from experimental images or data and is provided in the main text or figures. The reference cell radius $a_0$ was chosen such that the predicted cell volumes at the spheroid boundary are in good agreement with our experimental observations. From this analysis, we estimate the radii of CT26 and MDA-MB-231 cells to be 4.8 μm and 8.6 μm, respectively, values that align with those reported in the literature [12]. For calculations in **Figures 2, 3**, and **6**, we used 7.5 μm as a representative reference cell radius. The thickness of the actomyosin cortex in the cells is on the order of several hundred nanometers. For example, confocal imaging of F-actin in suspended MDA-MB-231 cells found an average cortex thickness of approximately 0.5 μm [12], which is the value used in our calculations. In **Figures 3C and 3D**, we quantify solid stress using the area-averaged magnitude of the stress, consistent with Ref. [13], defined as $< |\sigma_j| >= 1/A_s \int_{A_s} |\sigma_j|\, dA,\ j \in \{r, \theta\}$, where $A_s$ is the spheroid cross-sectional area over which the averaging is performed.

Living cells maintain an active cortical tension due to myosin II motor activity, which can be quantified through mechanical measurements. Using an AFM-based cell confinement assay, Hosseini et al. [14] measured the active contractile tension in the cortex of human breast, lung, and prostate cells. In MDA-MB-231 cells, this tension is approximately 0.33 mN/m. When converted to active in-plane stress by dividing by the ~0.5 μm cortex thickness, this corresponds to 0.66 kPa of contractile stress in the cortical layer. This value has been used in our calculations. Furthermore, reported values for the cortex Young's Modulus $E$ vary widely (0.2 – 30 kPa) for different cancer cells [15]. In this study, we assume values of $E = 9.6\ kPa$ for MDA-MB-231 cells based on experimental measurements [16], and $E = 6\ kPa$ for CT26 cells as a typical value [17]. Additionally, cell mechanical models commonly describe the cortical actin as a contractile, incompressible material [18]. This assumption aligns with direct measurements of Poisson's ratio in cancer (HeLa) cells [19], which show that the cortex is nearly incompressible over long timescales. Accordingly, we set $\nu = 0.5$ in our calculations.

**Table S1:** Model parameters and corresponding references.

| Parameter | Description | Value | Reference |
|---|---|---|---|
| $\boldsymbol{h}$ | Thickness of the cortex layer ($\mu m$) | 0.5 | [12] |
| $\boldsymbol{\sigma_{act}}$ | Active stress due to the myosin motors activity ($Pa$) | 660 | [14] |
| $\boldsymbol{\nu}$ | Poisson's ratio of the cortex | 0.5 | [18] |
| $\boldsymbol{g^+}$ | Ion channel conductance for positive ions ($S.m^{-2}$) | 0.1 | [20] |
| $\boldsymbol{g^-}$ | Ion channel conductance for negative ions ($S.m^{-2}$) | 2 | [20] |
| $\boldsymbol{x_{in}}$ | Intracellular impermeant species concentration (*mM*) | 100 | [21] |
| $\boldsymbol{z_{in}}$ | Average charge of the impermeant molecules | -1 | [20] |
| $\boldsymbol{c^+_{out}}$ | Concentration of positive ions outside the cell (*mM*) | 150 | [17] |
| $\boldsymbol{c^-_{out}}$ | Concentration of negative ions outside the cell (*mM*) | 110 | [17] |
| $\boldsymbol{x_{out}}$ | Extracellular impermeant species concentration (*mM*) | 40 | See Note |
| $\boldsymbol{p^+_0}$ | Physiological Na/K pump rate ($A \cdot m^{-2}$) | $21 \times 10^{-4}$ | See Note |
| $\boldsymbol{j}$ | Uniform oxygen consumption rate | See Note | See Note |
| $\boldsymbol{P_o}$ | Oxygen partial pressure at the spheroid boundary (mmHg) | 72 | [22] |
| $\boldsymbol{D}$ | Oxygen diffusion coefficient ($m^2/s$) | $2 \times 10^{-9}$ | [23] |
| $\boldsymbol{\Omega}$ | Constant (mmHg kg m$^{-3}$) | $3.0318 \times 10^7$ | [23] |
| $\boldsymbol{\rho_e}$ | Effective early proliferating fraction | 0.4 | See Note |
| $\boldsymbol{\rho_l}$ | Effective late proliferating fraction at the boundary | See Note | See Note |
| $\boldsymbol{w/r_{out}}$ | Normalized peripheral proliferation decay length | 0.15 | See Note |
| $\boldsymbol{\tau}$ | Spheroid surface tension (*mN/m*) | 6 | [24] |

Motivated by ex vivo measurements of tumor slices [25, 26] showing that extracellular ion concentrations remain unchanged across the tissue before widespread cell lysis, we assume that the external positive and negative ion concentrations remain at physiological levels across the tumor spheroid, with $c_{out}^{+} = 150\, mM$ and $c_{out}^{-} = 110\, mM$ [17]. As a result, to maintain electroneutrality in the interstitial space, the concentration of external negatively charged impermeant species in the spheroid, such as glycosaminoglycans, is estimated to be $x_{out} = 40\, mM$, assuming $z_{out} = -1$. We further assume that the conductivity coefficients are confined to previously suggested ranges $g^{+} = 0.1\, S.m^{2}$ and $g^{-} = 2\, S.m^{2}$ [20]. Additionally, experiments (often using X-ray microanalysis) have quantified the major inorganic ions inside cancer cells, from which the intracellular impermeant anion concentration ($x_{in}$) can be inferred. Across many types of cancer, the intracellular concentration of impermeant anions is on the order of 10² mM [21]. For example, human PC-3 prostate cancer cells have ~93 mM in impermeant anions, and HeLa cervical carcinoma cells ~97 mM. Accordingly, we set $x_{in} = 100\, mM$ in our calculations. Finally, based on the specified electrophysiological parameters and **Eqs. (7)–(10)**, we estimate the physiological sodium–potassium pump rate, $p_0^{+}$, in tumor spheroids to be $21 \times 10^{-4}\, A \cdot m^{-2}$ when the transmembrane hydrostatic pressure difference, $\Delta P$, is on the order of tens to a few hundred pascals (10–300 Pa).

Finally, we estimated oxygen consumption rates ($j$) and proliferative stretch parameters ($\rho_e$, $\rho_l$**,** and $w$) by fitting our model predictions of cell area and aspect ratio to the corresponding experimental data (**Figures 4** and **5**). The results indicate that the consumption rate of CT26 cells ($\approx 3.64 \times 10^{-6}\, \mathrm{m^3.kg^{-1}.s^{-1}}$) is higher than that of MDA-MB-231 cells, both in 11-day-old spheroids ($\approx 1.75 \times 10^{-6}\, \mathrm{m^3.kg^{-1}.s^{-1}}$) and in 14-day-old spheroids ($\approx 1.33 \times 10^{-6}\, \mathrm{m^3.kg^{-1}.s^{-1}}$). This observation is consistent with previously reported experimental measurements showing that the basal oxygen consumption rate of CT26 cells [27] is higher than that of MDA-MB-231 cells [28]. Furthermore, our estimated values fall within the range of oxygen consumption rates reported for other tumor spheroids [23]. In addition, our best fit to experimental data on partial oxygen pressure in human colon carcinoma spheroids [22] (**Figure 2D**) yields a consumption rate of $\approx 0.9 \times 10^{-6}\, \mathrm{m^3.kg^{-1}.s^{-1}}$, providing further validation of the physiological relevance of these estimates. Our results also indicate that the best fits are obtained with proliferative stretch parameters of $\rho_e = 0.4$, $w = 0.15 r_{out}$, and $\rho_l = 0.1$ for CT26 spheroids, and $\rho_l = 0.06$ for MDA-MB-231 spheroids.

**Supplementary Note S6:** *Residual solid stress field and cell morphology changes after core cell lysis*

When cells in the necrotic core of a tumor spheroid undergo lysis, they release a range of macromolecular debris [29, 30], many of which are negatively charged. These macromolecules span a broad size range, from nanometers to microns, which strongly influences their ability to diffuse through the extracellular pores of the tumor tissue. Experimental studies [31-33] suggest that pore sizes in the spheroid core typically range from 15 to 50 nanometers under physiological conditions. Consequently, large debris components, such as sub-micron DNA fragments [34], are unlikely to diffuse freely and tend to become trapped unless enzymatically degraded. We denote the concentration of these trapped negatively charged species as $x_{ly}$. To satisfy electroneutrality within the lysed core, the following condition must hold (**Eq. (10)**):

$$c_{ly}^{+} - c_{ly}^{-} - x_{ly} = 0 \tag{S27}$$

where the subscript "$ly$" refers to the necrotic core following widespread lysis of its constituent cells. Furthermore, passive diffusion of positive and negative ions between the lysed core and the surrounding interstitial spaces leads to an unequal distribution of these ions, which can be expressed as:

$$c_{ly}^{+} c_{ly}^{-} = c_{out}^{+} c_{out}^{-} \tag{S28}$$

This condition is known as the Donnan equilibrium. This result can be derived from **Eqs. (8)** and **(9)** by setting $p^+ = 0$ and assuming quasi-static conditions. Based on these relations, the hydrostatic pressure difference between the lysed core and the surrounding extracellular space (**Figure 5E**) can be expressed as:

$$\Delta P_{ly} = P_{ly} - P_{out} = \Delta\Pi_{ly} = RT(c_{ly}^+ + c_{ly}^- + x_{ly} - (c_{out}^+ + c_{out}^- + x_{out})) \tag{S29}$$

Utilizing **Eqs. (S27)–(S29)**, this pressure difference can be expressed as:

$$\Delta P_{ly} = RT(x_{ly} + \sqrt{4c_{out}^+ c_{out}^- + x_{ly}^2} - (c_{out}^+ + c_{out}^- + x_{out})) \tag{S30}$$

This expression is known as the Donnan (osmotic) pressure, and it governs the residual radial solid stress at the core boundary ($r = r_n$):

$$\sigma_r(r_n) = -\Delta P_{ly} \tag{S31}$$

As a result, our mechano-electro-osmotic (MEO) model can be extended to describe tumor spheroids with widespread core cell lysis by treating them as hollow spheroids with a specified radial stress at the inner radius (**Eq.** (**S31**)). All other governing equations of the MEO model remain unchanged and are valid for the region $r_n \le r \le r_o$. The value of $x_{ly}$ can be estimated from best fits to measured changes in cell area and aspect ratio across the spheroid. For the Day-14 MDA-MB-231 spheroid shown in **Figure 5C**, this estimate yields $x_{ly} \approx 40.18\, mM$.

**Supplementary Note S7:** *Spatial distributions of nuclear envelope tension and curvature predict rupture sites in elongated nuclei.*

Immunohistochemical analysis of advanced in situ human breast tumors [35] shows that extreme cellular elongation at the tumor periphery can compromise nuclear integrity, leading to rupture of the nuclear envelope (**Figure 6E**). Our model provides an explanation for this rupture by revealing how the primary driving forces, namely nuclear envelope areal stretch (tension) and Gaussian curvature, vary spatially across the envelope (**Figure S4**). Gaussian curvature is particularly important, as it regulates the local dilution of the nuclear lamina [36], which provides structural support to the envelope. In line with experimental observations (**Figure S4**, inset) [35, 37], we model the elongated nuclei at the tumor periphery as oblate in shape (**Figure S4**). This allows us to apply our previously derived theoretical results for cell cortex deformation (**Eqs. (S6)** and **(S7)**). These results indicate that the envelope areal stretch, defined as $\lambda_n^A = \lambda_\varphi \lambda_s$, can be mathematically expressed as:

$$\lambda_n^A = Vr_n^{\frac{2}{3}} Ar_n^{-\frac{1}{3}} \sqrt{\frac{Ar_n^4 + \tan^2(\psi)}{Ar_n^2 + \tan^2(\psi)}} \tag{S32}$$

where $Ar_n$ and $Vr_n$ denote the nuclear aspect ratio and the normalized nuclear volume (relative to its physiological value), respectively, and $\psi$ is the polar angle, as indicated in **Figure S4**. Furthermore, the Gaussian curvature $K$ of the oblate-shaped nucleus can be expressed as:

$$Ke_0^2 = \frac{Ar_n^{\frac{4}{3}} Vr_n^{-\frac{2}{3}}}{[Ar_n^2 \cos^2(\psi) + \sin^2(\psi)]^2} \tag{S33}$$

where $e_0$ is the nuclear radius under physiological conditions. Equations **(S32)** and (**S33)** show opposing spatial distributions of nuclear envelope areal stretch and Gaussian curvature with respect to the polar angle $\psi$ (**Figure S4**). Specifically, Gaussian curvature is maximal at the nuclear equator ($\psi = \pi/2$) and decreases toward the poles ($\psi = 0, \pi$), whereas envelope areal stretch (tension) is greatest at the poles and diminishes toward the equator. Consequently, nuclear rupture is predicted to occur between the equator and the poles, closer to the equator (pink region in **Figure S4**), where the nuclear envelope is structurally weaker and membrane tension remains sufficiently elevated. This prediction is consistent with experimental observations in mouse tumor xenografts (see [35] and the inset in **Figure S4**).

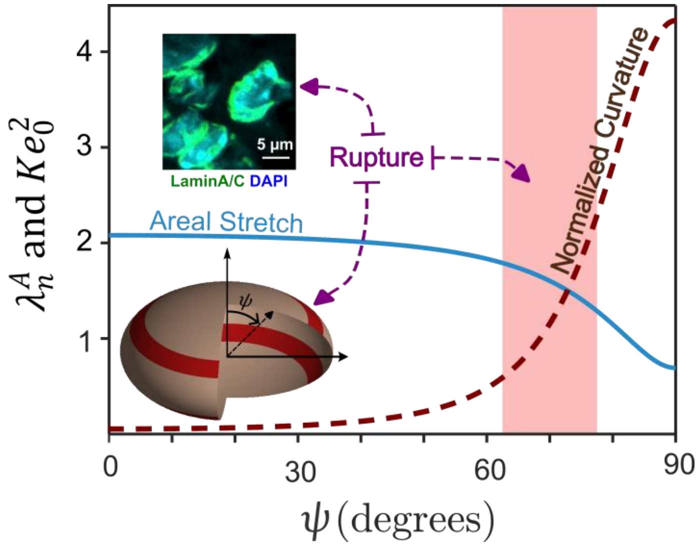


**Figure S4**: Spatial variation of nuclear envelope areal stretch (blue) and normalized Gaussian curvature (dark red) for an oblate nucleus with $Ar_n = 3$ and $Vr_n = 1$, showing predicted rupture between the equator and poles (pink region). Inset: immunohistochemical image of nuclear rupture at the tumor periphery from [35] (Lamin A/C in green, DAPI in blue).

**Supplementary Note S8:** *Published Smart-seq3D spheroid transcriptomics supports MEO predictions.*

Recently, diffusion-based dye labeling has been coupled to Smart-seq3 profiling to obtain deep single-cell transcriptomes together with a continuous core–periphery coordinate in triple-negative breast cancer spheroids [38]. In this dataset, each cell is assigned a radial position by log-transforming Calcein-AM fluorescence and rescaling it to the unit interval, such that $r = 0$ corresponds to the spheroid core and $r = 1$ to the boundary. We used this dataset in a preliminary analysis to evaluate whether transcriptional programs in large spheroids are consistent with key predictions of our mechano-electro-osmotic (MEO) model. This dataset is particularly suitable because it profiles the same cell line (MDA-MB-231) used in our in vitro experiments (Section 3.5) and employs large spheroids (diameter > 300 µm) [38], in which diffusion limitations are expected to generate pronounced core-to-periphery gradients. We first checked whether hypoxia-associated transcription is enriched toward the core, as expected for large spheroids. Using the MSigDB Hallmark Hypoxia gene set, we observed that core-like cells had higher hypoxia abundance than rim-like cells (**Figure S5A**; bottom 20% vs top 20% of $r$; Wilcoxon *p=0.00678*), consistent with the presence of a transcriptionally hypoxic inner compartment in these large spheroids [38].

We then tested the model's central mechanical prediction: moving from core to periphery, the solid-stress state transitions from compression-dominant to tension-dominant. We therefore asked whether a tensile-force–responsive transcriptional signature increases with radial position. We defined a conservative tensile-responsive gene set a priori from an independent mechanobiology RNA-seq dataset [39] in which breast cancer cells were exposed to graded cyclic tensile strain, selecting genes that increased strictly monotonically with strain magnitude (listed in **Table S2**). When quantified across the continuous radial axis in Smart-seq3D, this tensile-responsive program showed a clear periphery-increasing monotone trend (**Figure S5B**), supported by negative binomial regression with a quadratic dependence on $r$ (LRT $p = 2.67 \times 10^{-18}$) with an estimated periphery-to-core fold change (FC) of 2.07.

To further test mechanotransduction pathways expected to activate under increased tension and elongation, we next examined YAP/TAZ target programs. Using the MSigDB CORDENONSI_YAP_CONSERVED_SIGNATURE gene set, we found that YAP/TAZ target activity showed a significant non-linear spatial pattern, peaking at intermediate-to-outer radii and remaining modestly elevated at the periphery relative to the core (**Figure S5C***;* LRT *p=0.00976*; periphery/core FC = 1.23). Because broad YAP/TAZ signatures can include proliferation/cell-cycle–associated genes, we repeated the analysis using a smaller curated mini-set of canonical YAP targets *(*listed in **Table S3***)* to reduce potential cell-cycle confounding. This canonical YAP mini-set again peaked nearer the periphery and remained strongly elevated at the periphery relative to the core (**Figure S5D;** LRT $p = 2.41 \times 10^{-5}$*;* periphery/core FC = 2.26). Together, the hypoxia (**Figure S5A**), tensile-response (**Figure S5B**), and YAP/TAZ (**Figure S5C–D**) patterns provide complementary transcriptional support for the MEO model's predicted core–periphery transition from a compressed, hypoxic interior to a more tension-dominant, well-oxygenated peripheral state.

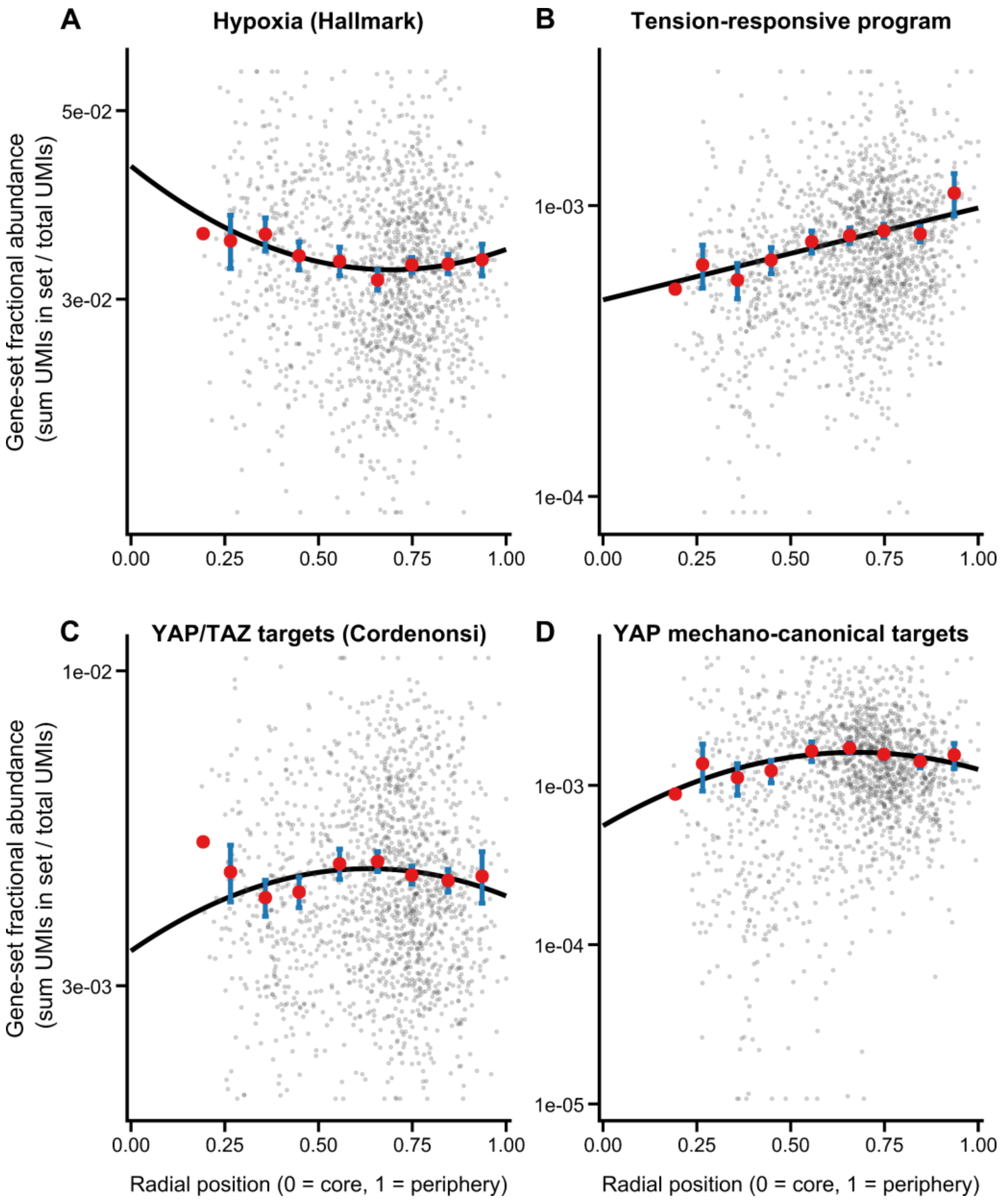


**Figure S5**: **Reanalysis of published diffusion Smart-seq3D spheroid transcriptomes reveals core-enriched hypoxia and periphery-enriched mechanoresponsive programs.** Data are from the diffusion Smart-seq3D study of MDA-MB-231 spheroids [38], in which each cell is assigned a continuous radial position $r$ (0 = core, 1 = periphery) derived from Calcein-AM fluorescence. In all panels, gene-set activity is quantified as gene-set fractional abundance (sum of UMIs in the gene set divided by total UMIs per cell). Grey points denote single-cell values, red circles indicate binned means (10 radial bins) with error bars reflecting within-bin variability, and black curves show fitted negative binomial trends across the full radial axis. **(A)** Hypoxia program activity (MSigDB *HALLMARK_HYPOXIA*) plotted against $r$; showing higher activity in core-like cells. **(B)** Tensile-force–responsive gene-set activity plotted against $r$, exhibiting a monotone increase toward the periphery, consistent with enhanced tensile-associated transcriptional programs outside the core. **(C)** YAP/TAZ target program activity using the MSigDB *CORDENONSI_YAP_CONSERVED_SIGNATURE*, displaying a non-linear spatial pattern with a peak at intermediate-to-outer radial positions and elevated activity at the periphery relative to the core. **(D)** Same analysis for a curated mini-set of canonical YAP targets to reduce potential cell-cycle/proliferation contributions. This canonical YAP program again peaks nearer the periphery and remains strongly elevated in peripheral cells.

**Table S2: Tensile-responsive gene set:** Defined a priori from an independent mechanobiology RNA-seq dataset [39] by selecting genes that increased strictly monotonically with increasing cyclic tensile strain magnitude**.**

| *Gene symbol* | *Notes* |
|---|---|
| *AKR1B1* | *Strict monotone increase with tensile strain* |
| *AKR1C1* | *Strict monotone increase with tensile strain* |
| *AIM1* | *Not detected in Smart-seq3D dataset; excluded from scoring* |
| *CYR61* | *Strict monotone increase with tensile strain* |
| *DLC1* | *Strict monotone increase with tensile strain* |
| *DUSP1* | *Strict monotone increase with tensile strain* |
| *ECT2* | *Strict monotone increase with tensile strain* |
| *GADD45B* | *Strict monotone increase with tensile strain* |
| *GLS* | *Strict monotone increase with tensile strain* |
| *SCHIP1* | *Strict monotone increase with tensile strain* |
| *SERPINE1* | *Strict monotone increase with tensile strain* |
| *SHCBP1* | *Strict monotone increase with tensile strain* |
| *TNS1* | *Strict monotone increase with tensile strain]* |
| *CD274* | *Strict monotone increase with tensile strain* |
| *CDH4* | *Not detected in Smart-seq3D dataset; excluded from scoring* |
| *CLDN1* | *Strict monotone increase with tensile strain* |
| *NEGR1* | *Strict monotone increase with tensile strain* |
| *PVR* | *Strict monotone increase with tensile strain* |

**Table S3:** Canonical YAP target mini-set: Curated a priori as a compact set of canonical YAP/TAZ-responsive targets**.**

| *Gene symbol* | *Rationale for inclusion* |
|---|---|
| *CYR61* | *Canonical YAP/TAZ target gene* |
| *CTGF* | *Canonical YAP/TAZ target gene* |
| *ANKRD1* | *Canonical YAP/TAZ target gene* |
| *AMOTL2* | *YAP/TAZ-regulated target gene* |
| *AREG* | *Canonical YAP/TAZ-responsive target gene* |
| *AXL* | *YAP/TAZ-regulated receptor tyrosine kinase* |
| *SERPINE1* | *YAP/TAZ-regulated extracellular matrix-associated gene* |

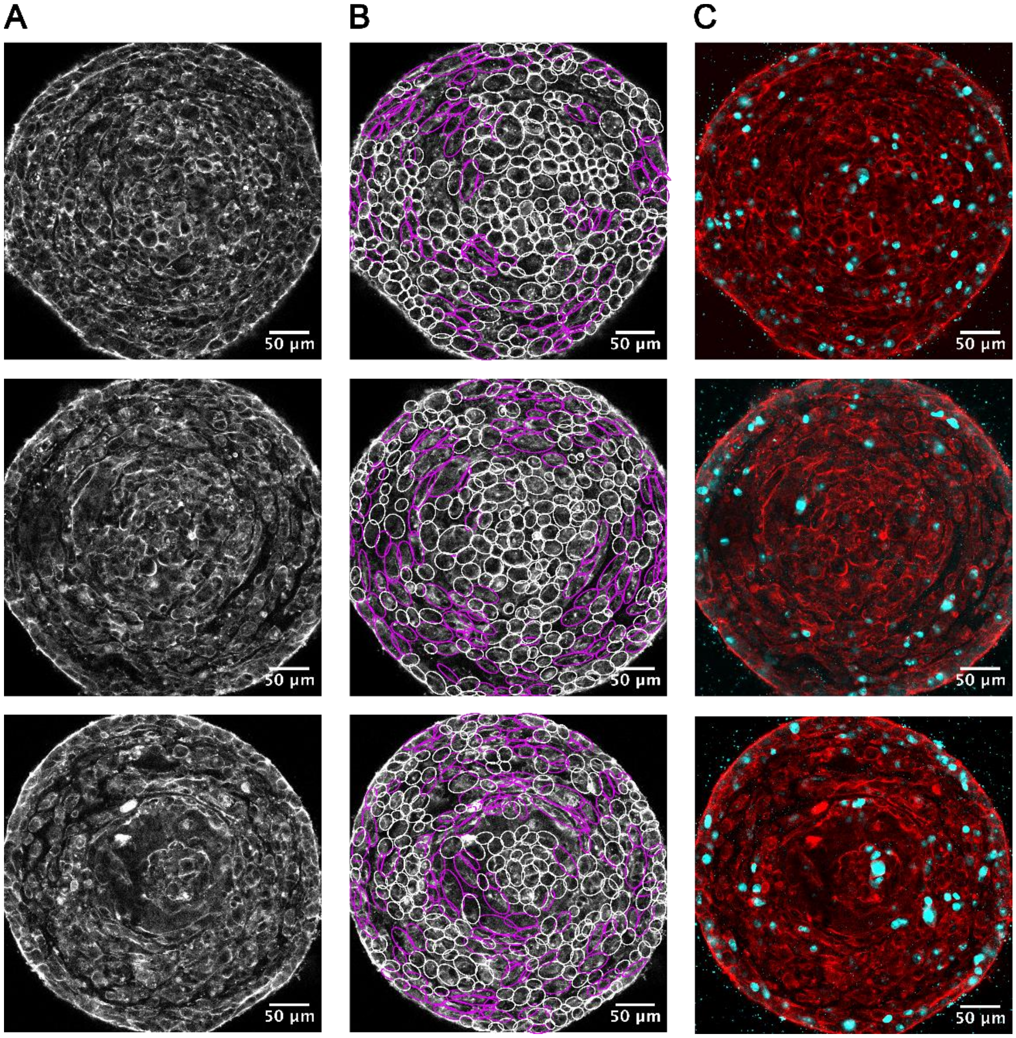


**Figure S6: Confocal mid-plane sections of 11-day-old MDA-MB-231 tumor spheroids.** **(A)** F-actin staining (phalloidin; shown in grayscale) highlights cortical structures and cell boundaries. **(B)** AI-based single-cell segmentation overlay showing cell boundaries (white) with elongated ellipse fits to visualize cell shape and orientation. Magenta outlines mark significantly elongated cells in the intermediate region (AR > 1.7). **(C)** F-actin (red) with Ki-67 (cyan) marking proliferating cells, enriched toward the spheroid periphery. Scale bars, 50 µm.

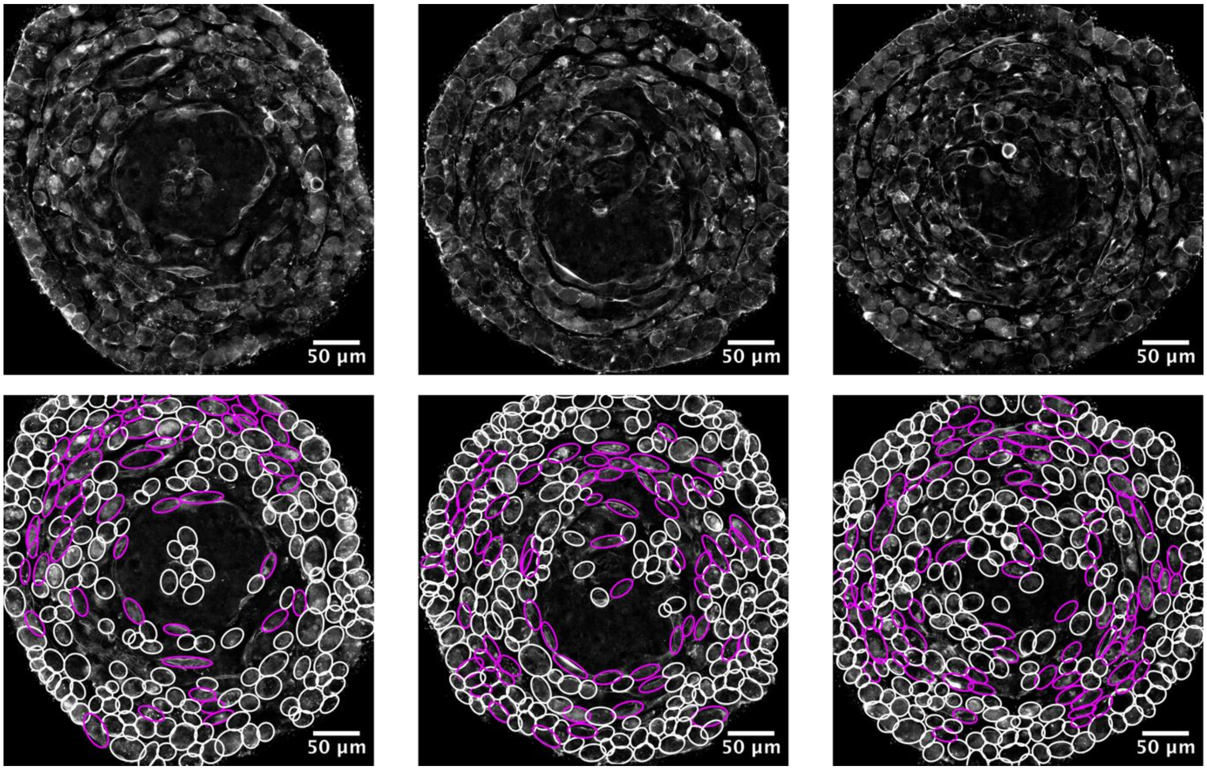


**Figure S7**: **Confocal mid-plane images of 14-day-old MDA-MB-231 tumor spheroids**. (Upper panel) Filamentous actin stained with phalloidin (shown in grayscale) highlights cortical structures and cell boundaries. (Lower panel) AI-based cell segmentation overlay showing individual cell shapes and areas. Magenta outlines mark significantly elongated cells in the intermediate region (AR > 1.7). Scale bars, 50 µm.

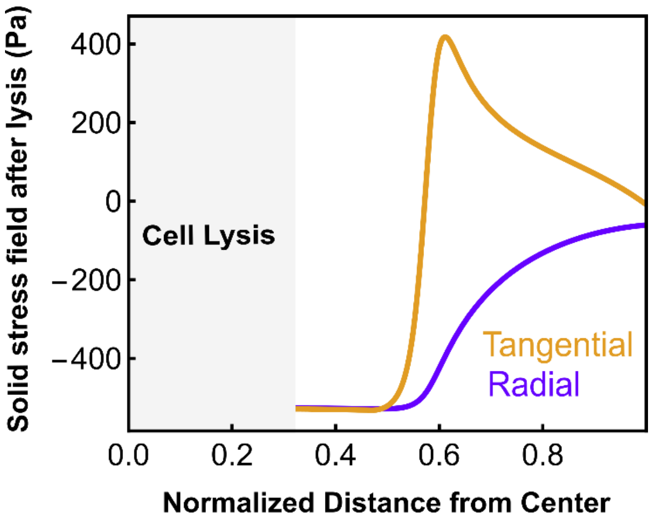


**Figure S8**: Model predictions of the solid stress field after core cell lysis in 14-day-old MDA-MB-231 tumor spheroids.

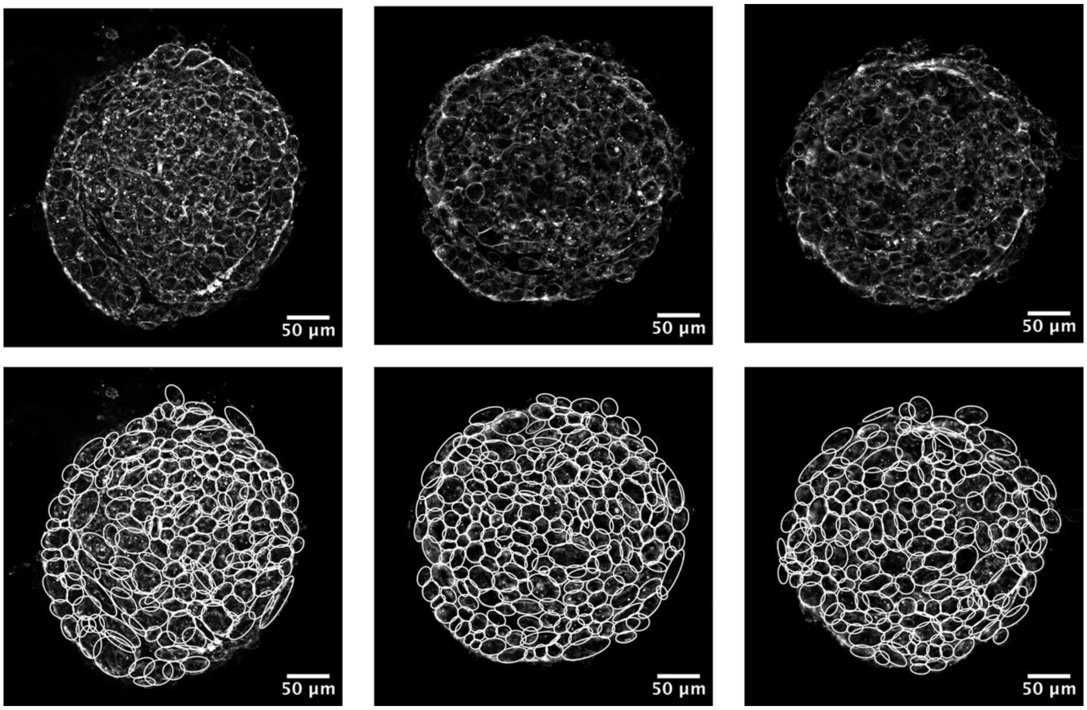


**Figure S9**: **Confocal mid-plane images of 4-day-old MDA-MB-231 tumor spheroids**. (Upper panel) Filamentous actin stained with phalloidin (shown in grayscale) highlights cortical structures and cell boundaries. (Lower panel) AI-based cell segmentation overlay showing individual cell shapes and areas. Scale bars, 50 µm.

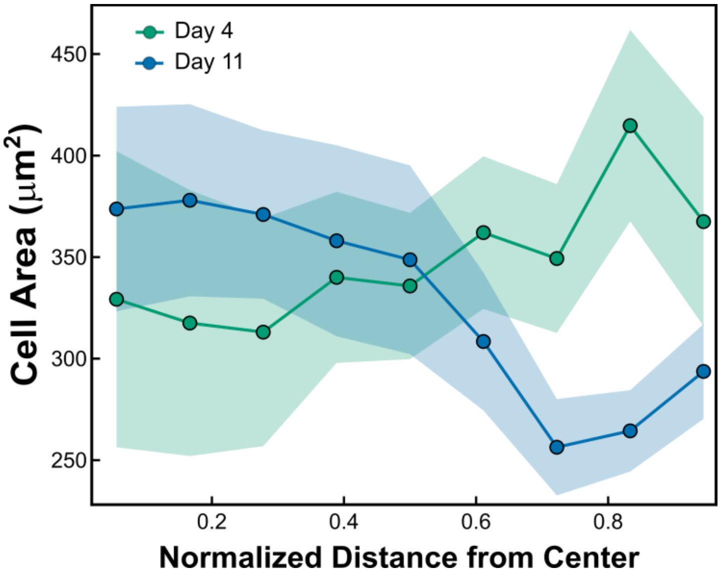


**Figure S10: Quantification of cell area as a function of normalized radial position for 4-day and 11-day spheroids.** Smaller spheroids show increasing cell area from core to periphery, whereas larger spheroids exhibit the reversed trend, with increased cell area in the core. Shaded regions denote ±SEM.

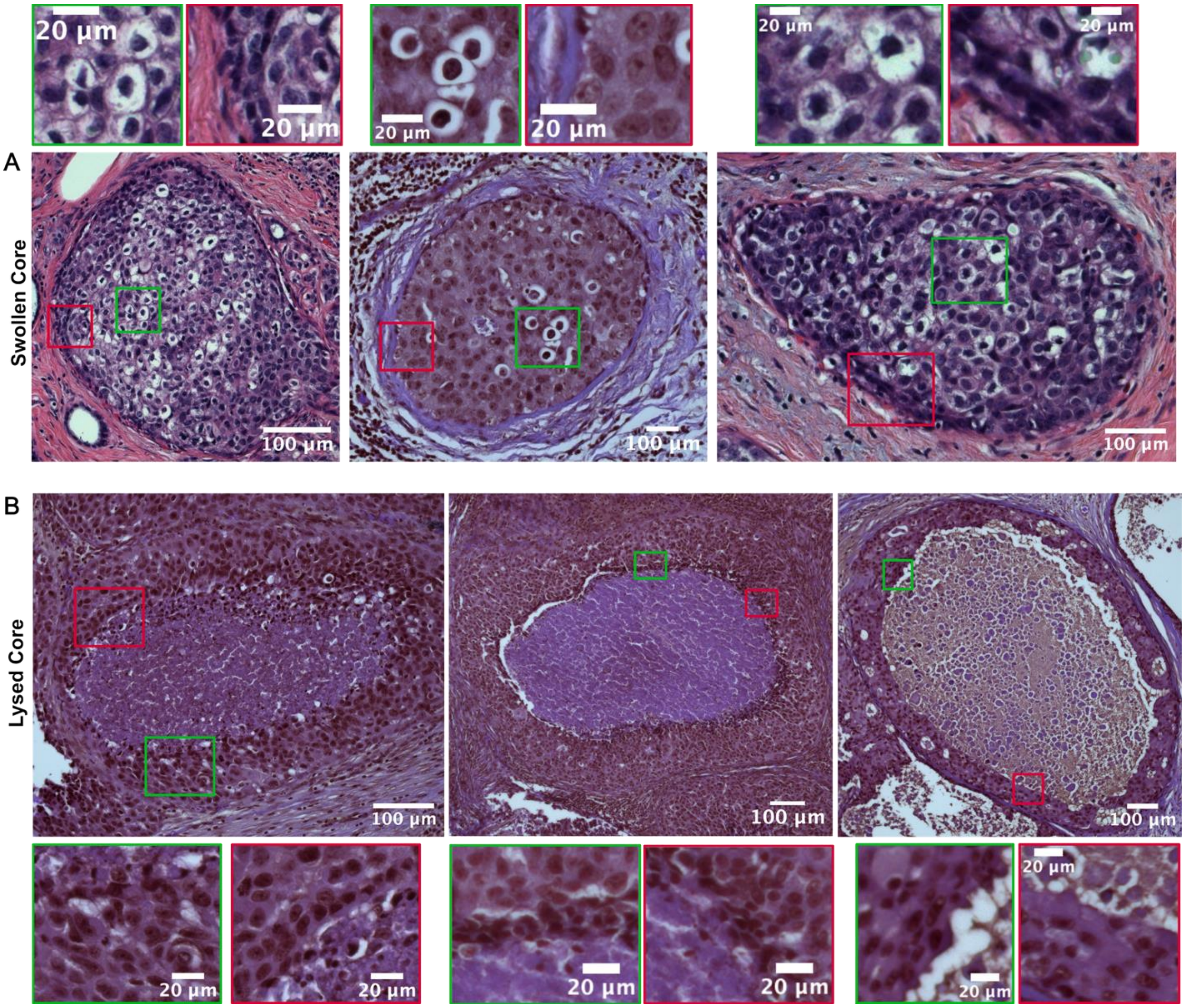


**Figure S11: Representative H&E-stained sections of human DCIS lesions illustrating swelling- and lysis-associated morphology. (A)** DCIS lesions without histologically visible necrosis show a swollen intraductal core with enlarged, rounded cells. Green boxes show magnified swollen cell regions and red boxes show magnified elongated cell regions. **(B)** DCIS lesions with central comedo necrosis show a lysed core surrounded by a viable rim of tumor cells. Green and red boxes show magnified elongated cell regions outside the lysed core. Scale bars, 100 µm (main panels) and 20 µm (insets).
